# A reference-quality NLRome for the hexaploid sweetpotato and diploid wild relatives

**DOI:** 10.1101/2025.01.13.632774

**Authors:** C. H. Parada-Rojas, K. L. Childs, M. Fernández de Soto, A. Salcedo, K. Pecota, G. C. Yencho, C. Almeyda, M. Kitavi, C. R. Buell, G. C. Conant, D. Baltzegar, L. M. Quesada-Ocampo

## Abstract

Breeding for sweetpotato (*Ipomea batatas*) resistance requires accelerating our understanding genomic of sources of resistance. Nucleotide-binding domain leucine-rich repeat receptors (NLRs) proteins represent a key component of the plant immune system that mediate plant immune responses. We cataloged the NLR diversity in 32 hexaploid sweetpotato genotypes and three diploid wild relatives using resistance gene enrichment sequencing (RenSeq) to capture and sequence full NLRs. A custom designed NLR bait-library enriched NLR genes with an average 97% target capture rate. We employed a curated database of cloned and functionally characterized NLRs to assign sequenced sweetpotato NLRs to canonical phylogenetic clades. We identified between 800 to 1,200 complete NLRs, highlighting the expanded diversity of coiled-coil NLRs (CNLs) across all genotypes. NLRs among sweetpotato genotypes exhibited large conservation across genotypes. Phylogenetic distance between 6X (hexaploid) and 2X (diploid) genotypes revealed that a small repertoire of *I. batatas* CNLs diverged from the sweetpotato wild relatives. Finally, we obtained chromosome coordinates in hexaploid (Beauregard) and diploid (*Ipomoea trifida*) genomes and recorded clustering of NLRs on chromosomes arms. Our study provides a catalog of NLR genes that can be used to accelerate breeding and increase our understanding of evolutionary dynamics of sweetpotato NLRs.

## INTRODUCTION

Opportunistic plant pathogens survive in agroecosystems strategically by widening their host range within and between plant families and evolving long term survival structures (Henry *et al*. 2019). Soilborne plant pathogens (nematodes, fungi, oomycetes, and bacteria) have evolved strategies to persist endemically and reemerge to infect susceptible hosts when the opportunity presents itself (Quesada-Ocampo *et al*. 2023). In the last decade, restrictions on soil fumigation resulted in the reemergence of many soilborne pathogens and pests in agriculture (Chellemi *et al*. 2016; Holmes *et al*. 2020; Land *et al*. 2022; Sanogo *et al*. 2022). Without effective soil fumigation, farmers rely on a combination of fungicides/nematicides, biological amendments, and host resistance (Miller *et al*. 2020; Parada-Rojas and Quesada-Ocampo, 2022). Host resistance remains a tool that provides flexible and economical control that is compatible with diverse cropping systems (Michelmore *et al*. 2017). However, breeding for pathogen resilient crops requires accelerating our understanding of sources of resistance within plant genomes. Many domesticated crop draft genomes have been sequenced and refined (Sato *et al*. 2012; Kim and Buell, 2015; Consortium (IWGSC) *et al*. 2018; Edger *et al*. 2019), yet the vast majority of crop species lack a comprehensive understanding of resistance loci within their genomes to realize their full potential (Steuernagel *et al*. 2015; Kourelis and van der Hoorn, 2018; Parada-Rojas and Quesada-Ocampo, 2021).

Plants and pathogens exist in a continuum of coevolutionary struggle for survival (Tamborski and Krasileva, 2020; Derevnina *et al*. 2021). An important layer of plant immunity is the recognition of pathogen effector molecules by innate intracellular immune receptors known as NLR proteins (nod-like receptors or nucleotide binding leucine rich repeat proteins) (Jones *et al*. 2016). NLR proteins are part of a larger class of cellular receptors used for chemical communication within and between organisms, and a subset of NLRs have evolved to stimulate defense response in cells (Steidele and Stam, 2021). Upon direct or indirect recognition of secreted pathogen effectors, NLRs induce robust immune responses that include among other features hypersensitive programmed cell death, which has the potential to render plants resistant (Balint-Kurti, 2019). NLR architecture includes a combination of a canonical nucleotide binding (NBARC) domain, a C-terminal leucine rich repeat (LRR) domain, and three diverse accessory N-terminal domains, Toll-interleukin-1 receptor (TIR), coiled-coil (CC), and Resistance to Powdery Mildew 8 (RPW8) domains. Generally, patterns of N-terminal domain composition inform the classification of NLRs into three major classes that form monophyletic groups in the NLR phylogeny: TNLs, CNLs, and RNLs (Shao *et al*. 2016; Tamborski and Krasileva, 2020; Kourelis *et al*. 2021). Other subgroups of NLRs include those that carry the late-blight R1 (B) domain, the recently discovered C-terminal jelly roll/Ig-like domain (C-JID/J), and any non canonical integrated domains (IDs) (Ballvora *et al*. 2002; Cesari *et al*. 2014; Ma *et al*. 2020; Kourelis *et al*. 2021). NLRs have evolved as one of the most diverse gene families in plants in response to the extraordinary diversity of plant pathogens and their arsenal of protein effectors (Wu *et al*. 2017; Adachi *et al*. 2019). Collective NLR repertoire (NLRome) evolution favors networking among NLRs with sensor NLRs detecting pathogen effectors and helper NLRs rendering effector recognition into a hypersensitive response (HR) phenotype (Derevnina *et al*. 2021). These two types of NLRs are considered key components of the NLR immune network and are more recently referred to as NLR-required for cell death (NRC) proteins. NRCs represent a phylogenetically supported NLR class with deployment potential (Wu *et al*. 2017; Kourelis *et al*. 2022).

Because of the sheer diversity in this plant gene family, genome assembly and annotation of NLRs in different plant species often requires novel approaches (Jupe *et al*. 2013; Stam *et al*. 2016; Witek *et al*. 2016; Van de Weyer *et al*. 2019). Even diploid whole genome projects struggle to generate accurate NLR annotations due to their clustering in chromosomes and overlap of repetitive sequences (Andolfo *et al*. 2014; Bayer *et al*. 2018). To add complexity, cultivated/domesticated polyploid crops often rely on knowledge of diploid wild relative genomes for crop improvement (Fajardo *et al*. 2016; Wu *et al*. 2018b; Edger *et al*. 2019; Sun *et al*. 2020). Target capture NLR sequencing (RenSeq) represents a desirable tool to reduce complexity and explore NLR diversity in crop species, especially if cultivated/domesticated genomes are available (Witek *et al*. 2016). One of the advantages of using exome capture tools is the ability of baits to hybridize to sequences with 20% mismatch (Witek *et al*. 2016; Giolai *et al*. 2016). This feature allows for the use of wild relative genomes to investigate complex domesticated polyploid crop species. RenSeq recently allowed examination of intra and interspecies NLR diversity in 64 *Arabidopsis thaliana* ecotypes and 16 accessions from 5 different *Solanum* species (Van de Weyer *et al*. 2019; Seong *et al*. 2020).

Sweetpotato (*Ipomoea batatas* (L.) Lam)(2n = 6X = 90), a globally grown root crop with origins from northern South America and Central America, provides more nutrients per farmed hectare than any other food crop (Oke *et al*. 1990; Truong *et al*. 2018). Due to its versatility as a staple feed, food, and fuel source, sweetpotatoes consistently rank highly among the most important crops worldwide (FAOSTAT, 2022). Like potato and numerous other flowering plants, cultivated sweetpotato exhibits polyploidy (6X) and high degrees of self-incompatibility (Arumuganathan and Earle, 1991; Tsuchiya, 2014). Its 1.6 Gb genome harbors a high degree of heterozygosity driven by outcrossing breeding methods (Wu *et al*. 2018b). Today, sweetpotato improvement through genomic selection mainly relies on high quality genomic resources of two diploid wild relatives (*I. trifida* and *I. triloba*) (da Silva Pereira *et al*. 2020; Oloka *et al*. 2021).

Identification and deployment of resistance in sweetpotato to emerging and persistent pathogen threats lags behind other staple crops (Chakraborty *et al*. 2018; Kaloshian and Teixeira, 2019; Wang *et al*. 2021). *Ceratocystis fimbriata*, *Fusarium solani* and *Meloidogyne enterolobii* represent contemporary soilborne pathogens that limit sweetpotato production and global trade (Lewthwaite *et al*. 2011; Scruggs and Quesada Ocampo, 2016; Yang *et al*. 2018; Lee *et al*. 2019; Schwarz *et al*. 2021; Parada-Rojas *et al*. 2021; Rutter *et al*. 2021). Host resistance represents a sustainable tool for sweetpotato resilience against pathogens that can protect cultivated sweetpotatoes globally. RenSeq is a novel genomic tool that can be used to reduce the complexity of the sweetpotato genome and reveal sweetpotato NLR gene diversity. A better understanding of the origin and evolution of NLR gene families in plants requires data from highly heterozygous polyploid domesticated staple crops. Here we describe in detail the first comprehensive NLRome in a non-model organism and a major crop species by cataloging 32 sweetpotato (6X) and 3 wild relative (2X) NLRomes. Specifically, we aimed to (i) compare the success of a RenSeq approach in recovering full NLR gene models in hexaploid *I. batatas*, and diploid *Ipomoea* species against whole genome sequencing annotations; (ii) dissect NLR domain diversity and evaluate their phylogenetic relationship among *I. batatas* and wild relatives; (iii) identify core and accessory NLRs in hexaploid sweetpotato based on NLR families; and (iv) provide a genetic map of NLRs in sweetpotato to help accelerate breeding efforts in sweetpotato. Our study provides a foundation for accelerating resistance breeding and functional studies of NLR genes in sweetpotato.

## RESULTS

### RenSeq sequencing and assembly quality

To catalog the NLR repertoire in sweetpotato, we implemented the long-read PacBio RenSeq protocol. Our RenSeq bait library was designed to capture 2,032 NLR coding regions with the bait library covering 90.1% of desired target positions with at least 1 bait. Library preparation and sequencing yielded a total of 6,163,025 CCS reads across all 35 genotypes with an average 176,086 reads per genotype. We consistently obtained above 90% on-target capture rate with an average of 97% of the CCS reads containing bait sequences at or above the set threshold across all 35 genotypes (Table S1). To evaluate read quality, we used NLR-parser’s definition of complete NLRs to calculate the number of reads containing complete NLR motifs for each sample. We found that on average 70% of the CCS reads carried a complete set of motifs associated with NLRs (Table S1). Combined CCS read metrics provided confidence in our bait library design and allowed us to accurately assemble NLRs in hexaploid sweetpotato. The captured and assembled NLRomes for hexaploid and diploid genotypes averaged 21.2 Mb and 7.8 Mb with contig N50 length of 6,208 and 7,372 bp, respectively (Table S2). The number of hexaploid RenSeq contigs ranged between 5,027 to 3,048 for hexaploid genotypes and 1,558 to 765 for diploid genotypes with an average of 3,556 and 1,145 contigs, respectively (Table S2). The resulting RenSeq assemblies for all genotypes ranged in coverage between 20X for Covington and 69X for Southern Delight with an average coverage across 35 genotypes of 42X. The NLR-annotator analysis revealed that on average 88% and 79% of the hexaploid and diploid genotype contigs contained NLR motifs, respectively. An average of 82% of the total number of NLR motif-containing contigs in both 6X and 2X genotypes carried NLRs defined as complete by NLR-annotator. On average and as defined by NLR-annotator, the number of contigs containing a single NLR complement for hexaploid and diploid genotypes was 2,795 and 790 respectively (Table S2).

**Table S1.** Enrichment quality assessment based on number and percentage of circular consensus reads (CCS) containing 1 or more target baits in a range of 96 base pairs at 80% sequence identity. This table also includes the number of reads identified by NLRparser as containing a complete or partial set of NLR motifs. *https://doi.org/10.6084/m9.figshare.21899886*

**Table S2.** Assembly statistics and NLR-Annotator based counts for 32 sweetpotato and 3 wild relative genotypes. *https://doi.org/10.6084/m9.figshare.21899898*

### RenSeq improves NLR annotation

Over the course of this study a sweetpotato hexaploid chromosome level assembly was released (http://sweetpotato.uga.edu). This prompted us to evaluate the performance of the standard annotation project versus our NLR tailored annotation pipeline. We expanded our search and included comparisons for plant species with completed RenSeq projects. These included mainly Solanales species and Arabidopsis. In the Beauregard 6X proteome and NLRome, we identified 2,471 and 2,871 proteins as NLRs, respectively (Figure 1). Sweetpotato ranked highest for NLR counts across all tested species. Overall, RenSeq projects annotated more NLRs than standard genome annotations. We found that *I. trifida* and *I. triloba* genome annotations consistently had lower counts in comparison to our RenSeq annotations. In particular, our *I. trifida* NLR tailored annotation yielded 3 times the NLR content of the genome project for the same genotype (Figure 1A). This analysis also revealed that NLR content is largely independent of genome size, with sweetpotato genome size (1.6Gb) being roughly half the pepper genome size (3.5Gb) but harboring more than double the number of NLRs (Figure 1B). These results indicate that RenSeq improves NLR annotation of the sweetpotato reference genome and its wild relatives.

**Figure 1.**
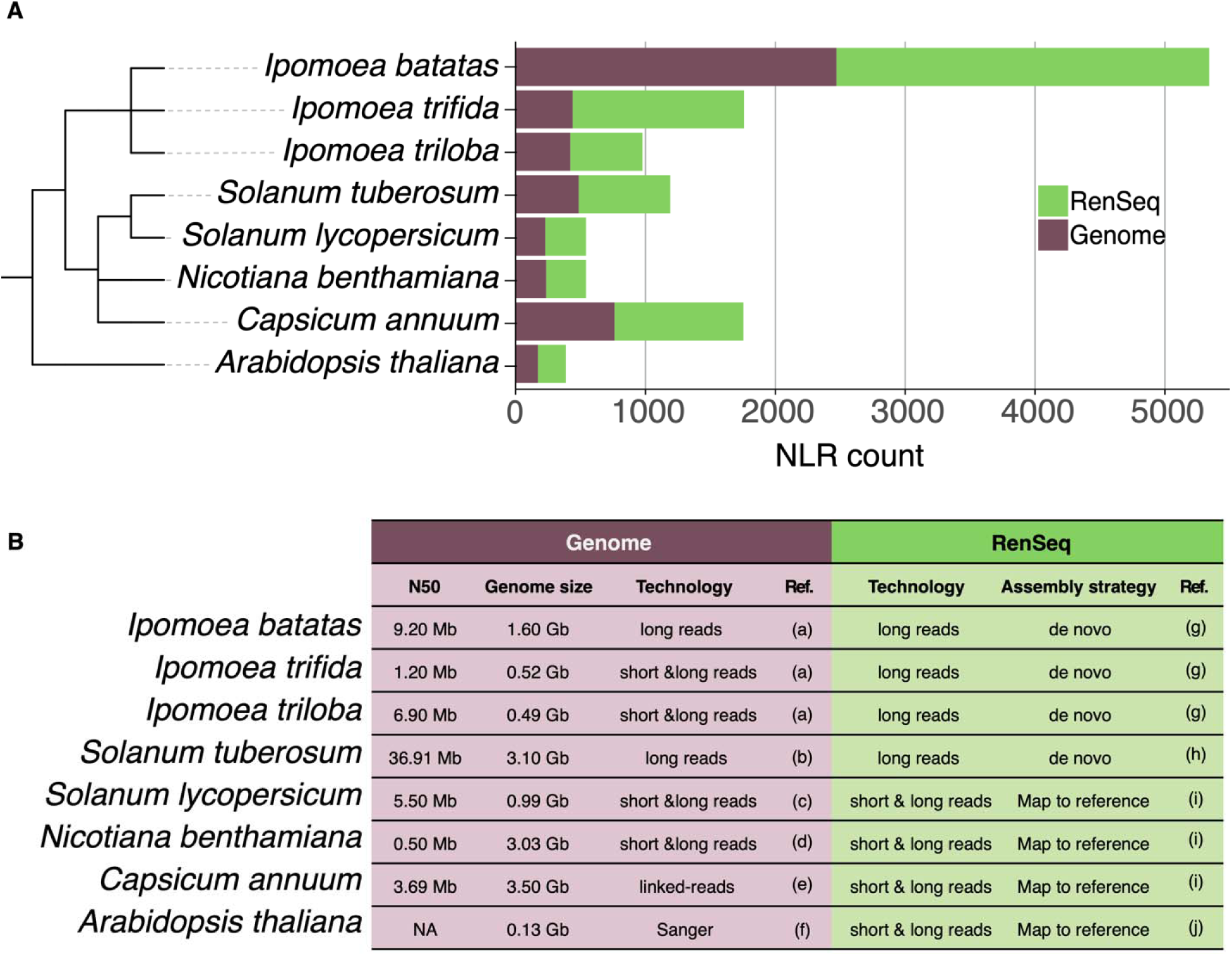
RenSeq improves NLR annotation. Species tree of a subset of Solanales species NLR annotations using RenSeq and Genome annotations. The numbers of nucleotide-binding and leucine-rich repeat immune receptors (NLRs) annotated per plant species as reported by each RenSeq effort versus the predicted annotation via NLRtracker from each proteome. (**A**) The species tree indicates the phylogenetic relationship of the species analyzed. The number of NLRs as annotated by NLRtracker is shown in the stack bar plot with green and brown bars representing RenSeq annotated and genome annotated NLRs for each species, respectively. (**B**) Genome statistics and sequencing technology used for both Genome and RenSeq projects. Refs. (a) Wu et al., 2018b, (b) Pham et al., 2020, (c) Hosmani et al., 2019, (d) Bombarely et al., 2012, (e) Hulse-Kemp et al., 2018, (f) TAIR, 2022, (g) This study, (h) Jupe et al. 2013, (i) Seong et al. 2020, (j) Van de Weyer et al. 2019.

### Sweetpotato and wild relative genomes harbor a diverse catalog of NLRs

To reveal the nature and the diversity of sweetpotato NLRs, we performed a comparative NLRome analysis that relies on NLR annotations by NLRtracker (Kourelis *et al*. 2021). Collectively, NLRtracker detected NLRs, degenerate NLRs, and NB-ARC containing proteins (Table S3). Informed by the NLRtracker results registered across all 35 genotypes, we arbitrarily focused on 13 main NLR domain architectures that we grouped into 5 major domains including the canonical CNL, RNL and TNL but also the non canonical NLR-IDs and BNLs (Figure S1). We categorized CNLs as NLRs containing one or two CC domains at the N-terminal and containing the NB-ARC and LRR domains. TNLs included NLRs containing TIR, NB-ARC and LRR domains in addition to an optional CID-J domain at the C-terminal. RNLs were categorized as NLRs containing RPW8, NB-ARC, and LRR domains. We also included the BNL category reported by NLRtracker, as NLRs containing a late blight R1 domain in the N-terminus, combined with CC, NB-ARC, and LRR domain (i.e. BNLs and BCNLs). We included the NLR Integrated Domains (NLR-IDs) as they represented a substantial proportion of NLRs found across all genotypes and potential effector targets. These NLRs were classified as NLR-IDs if they carried an ID (O) domain at either end of a complete NLR protein with the exception of ONLs, which could represent a novel N-terminal domain for sweetpotato. We excluded from our analysis NLRs that carried only an NB-ARC and a LRR domain as we consider them not full-length NLRs.

**Table S3.** NLRtracker output for each catalog of annotated NLRs in each of the 32 sweetpotato and 3 wild relative genotypes. *https://doi.org/10.6084/m9.figshare.21899910*

Characterization of the five NLR domains revealed a diverse set of architectures with a notable expansion of the CNLs across the 35 genotypes followed by TNLs and NLR-IDs (Figure S2). RNLs and BNLs represented a small fraction of the NLRs across all genotypes investigated (Figure 2). The wild relatives contained a proportionally similar number of CNLs, ranging from 111 to 272 CNLs. Among the *I. batatas* genotypes, Regal ranked highest for total NLR content followed by the historical genotype Apache (Figure 2). NC07-0847 and Centennial genotypes contained the lowest NLR counts across all *I. batatas* genotypes. We observed a consistent proportion of NLR-IDs across all *I. batatas* and wild relative genotypes as the third most-common NLR architecture we recorded in our study. Altogether, our NLR domain architecture analysis confirms the widespread presence of CNLs, TNLs, and NLR-IDs among sweetpotato genotypes.

**Figure 2.**
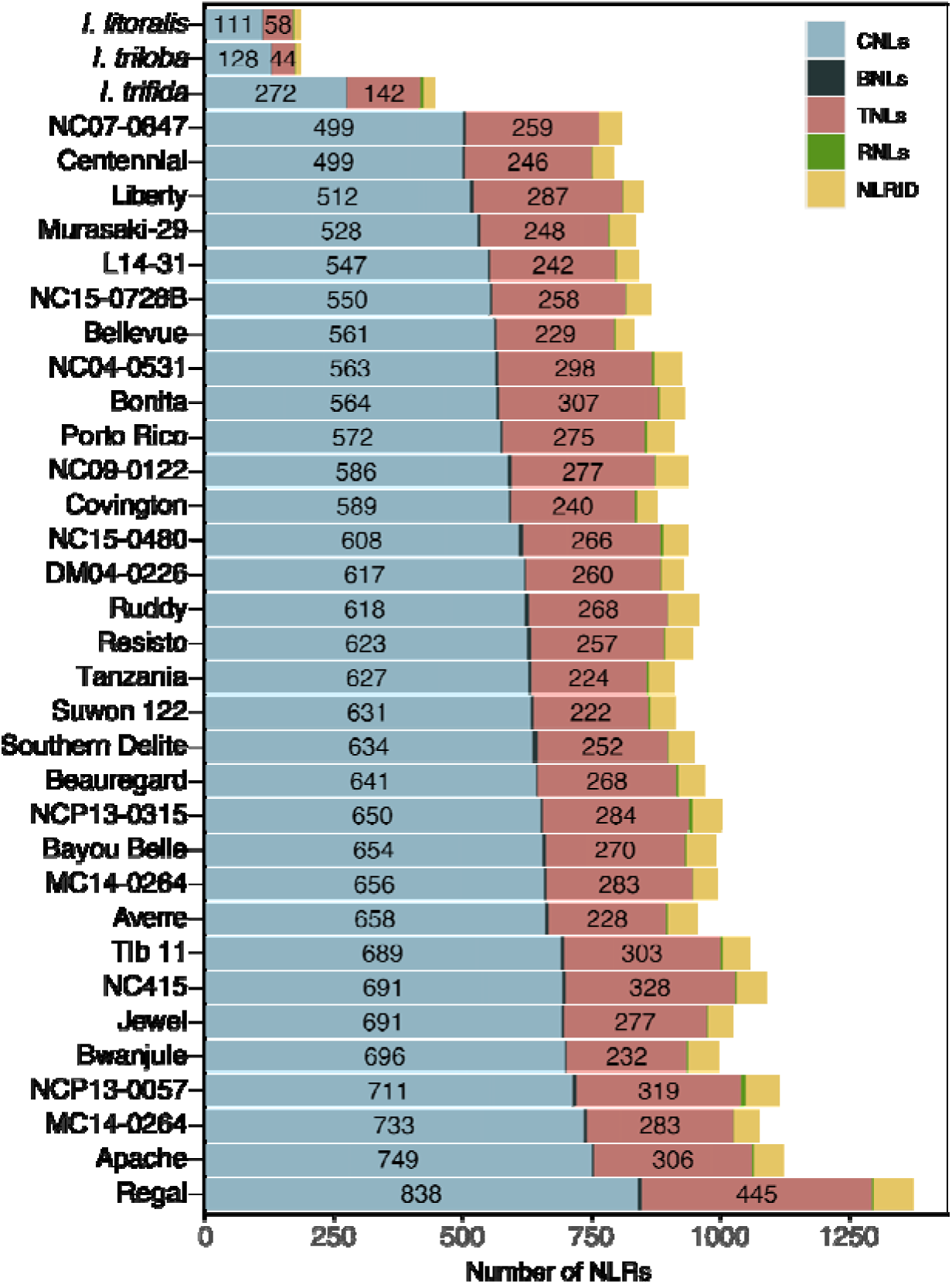
Sweetpotato and wild relative genomes harbor a diverse catalog of NLRs. Stack bar plot distribution of 32 sweetpotato genotypes and three *Ipomoea* spp. complete NLRs as annotated by NLRtracker. The number of each domain architecture for each genotype is plotted as a stack plot. CNLs, coiled-coil nucleotide-binding and leucine-rich repeat immune receptors (i.e. CNL or CCNL); BNLs, Late-Blight R1 nucleotide-binding and leucine-rich repeat immune receptors (i.e. BNL or BCNL); TNLs, Toll/interleukin-1 receptor nucleotide-binding and leucine-rich repeat immune receptors with or without C-terminal jelly roll/Ig-like domain (i.e. TNL or TNLJ); RNLs, N-terminal RPW8-type coiled-coil nucleotide-binding and leucine-rich repeat immune receptors; NLR-IDs, nucleotide-binding and leucine-rich repeat immune receptors containing non canonical “integrated domains”. Detailed domain architecture and abbreviations are as shown in Figure S1.

To examine the evolutionary history of NLRs in sweetpotato and its wild relatives, we constructed an NLR phylogeny using the NB-ARC domain and inferred major phylogenetic clades. The un-rooted 29,553 NB-ARC phylogeny clustered by the three canonical NLR domains: CNL, TNL and RNLs (Figure S3). To navigate the phylogeny, we placed 35 RefPlantNLRs and rooted the tree at the TNL clade. The CNL clade represented the largest domain architecture, with strong radiation and branching pattern, indicating high diversification within the CNL clade (Figure 3). Approximately half of the CNL clade expands beyond the anchoring of the RefPlantNLRs in our tree. The TNL and RNL clades were more compact in comparison to the CNL clade. We observed clustering of the BNLs across all genotypes within the CNL clade (Figure 3). To take a closer look at NLR-IDs within our phylogenetic tree, we decorated the outer ring that corresponded to NB-ARCs associated with NLR-IDs classified by NLRtracker (Kourelis *et al*. 2021). We observed the spread of NLR-IDs across the phylogeny with certain clades harboring more NLR-IDs than others (Figure 3). We inspected the NLR-ID clustering by canonical architecture with CNLO and OCNLs placement occurring within the CNL clade and TNLO and OTNL placement within the TNL clade (Figure S4). Notably, ONLs were mainly placed within the CNL clade, with a small number of ONLs falling within the TNL clade (Figure S4). We observed two large CNL sub-clades that had poor NLR-ID assignments (Figure S4). Our phylogenetic analysis highlights the expanded diversity of CNLs across all genotypes and their evolutionary history.

**Figure S3.**
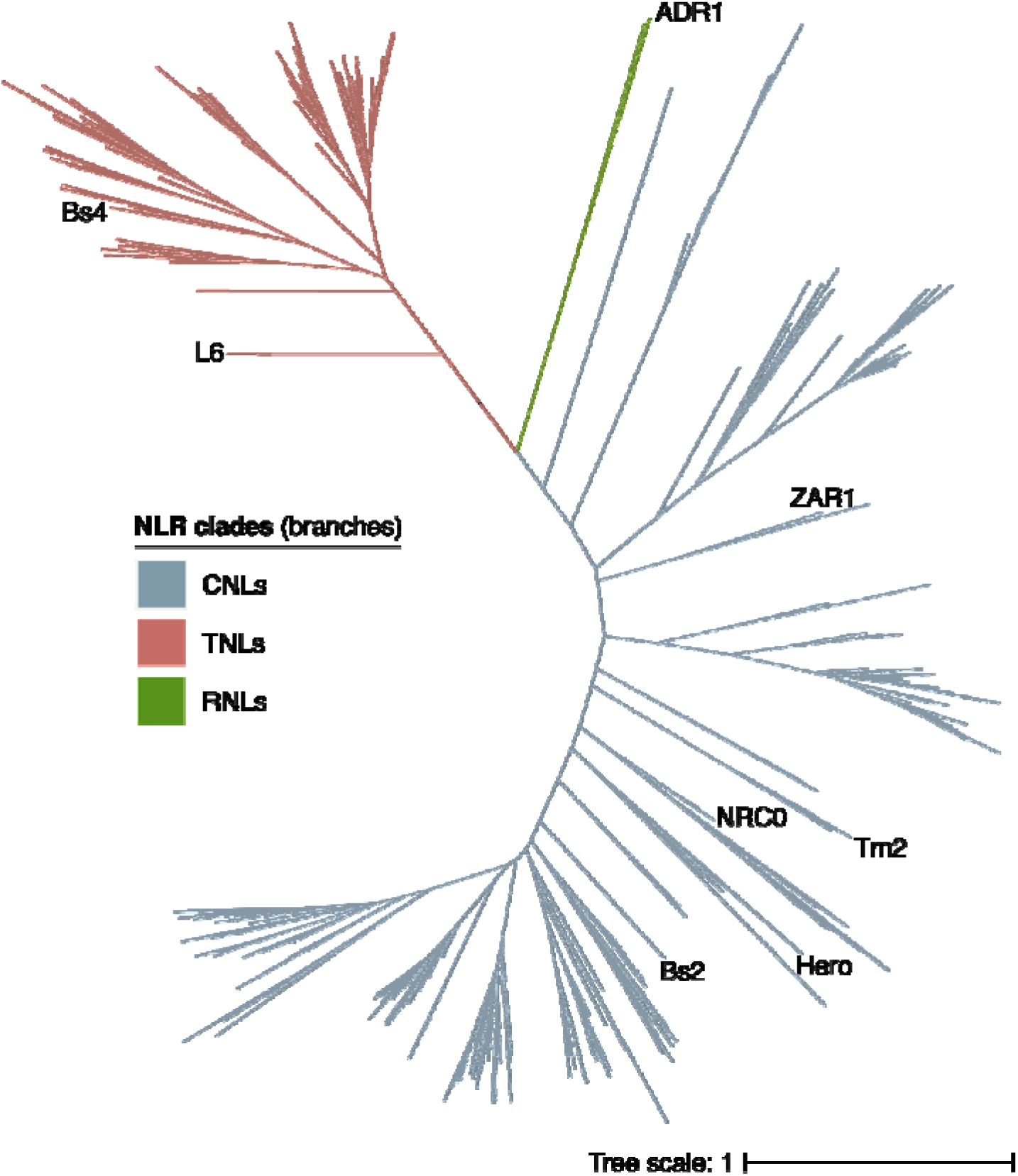
Unrooted sweetpotato and wild relatives NLR phylogeny exhibit clustering by canonical domain. Phylogenetic diversity of sweetpotato and wild relative NLRs. Unrooted NB ARC domain phylogeny of 29,553 amino acid sequences inferred using the Maximum Likelihood method based on the Jones–Taylor–Thornton (JTT) and Per Site Rate (PSR) models in ExaML. The major TNL, CNL, and RNL clades are indicated by branch colors. Tip labels correspond to RefPlantNLRs included in the phylogeny. Domain architecture and abbreviations are as shown in Figure S1. Branch scale represents the number of substitutions per site.

**Figure 3.**
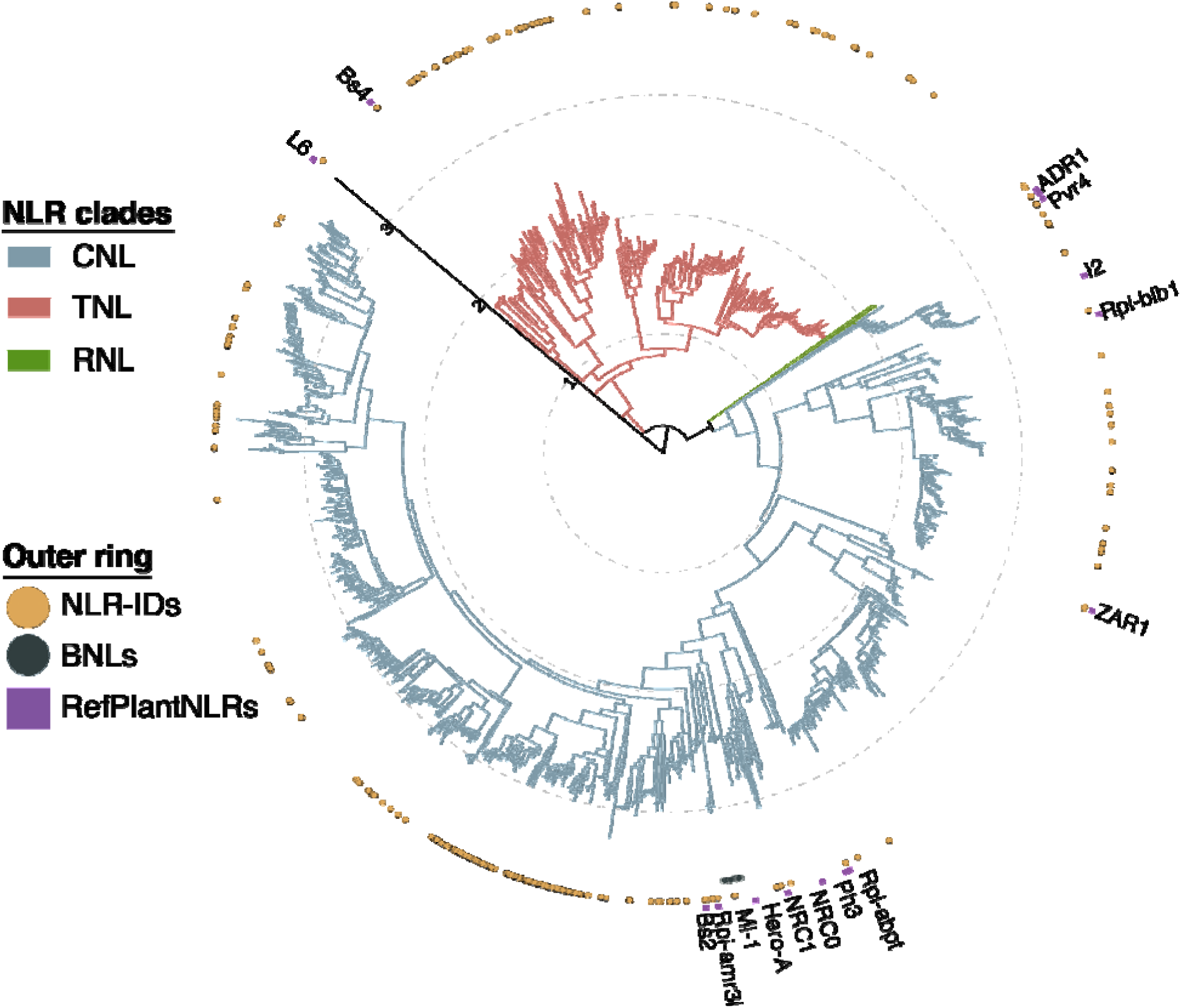
Sweetpotato and wild relatives exhibit expanded diversity of CNLs. Phylogenetic diversity of sweetpotato and wild relative NLRs. NB-ARC domain phylogeny of 29,553 amino acid sequences inferred using the Maximum Likelihood method based on the Jones Taylor Thornton (JTT) and Per Site Rate (PSR) models in ExaML. Domain architecture abbreviations correspond to CNLs, coiled-coil nucleotide-binding and leucine-rich repeat immune receptors (i.e. CNL or CCNL); BNLs, Late-Blight R1 nucleotide-binding and leucine-rich repeat immune receptors (i.e. BNL or BCNL); TNLs, Toll/interleukin-1 receptor nucleotide-binding and leucine-rich repeat immune receptors with or without C-terminal jelly roll/Ig-like domain (i.e. TNL or TNLJ); RNLs, N-terminal RPW8-type coiled-coil nucleotide-binding and leucine-rich repeat immune receptors; NLR-IDs, nucleotide-binding and leucine-rich repeat immune receptors containing non canonical “integrated domains” as shown in Figure S1. The tree branches are rooted on the branch connecting TNL and non-TNL clades. The major TNL, CNL, and RNL clades are indicated by branch colors. The color code of the outer ring shapes indicates to which of the non canonical NLR architectures (i.e. NLR-IDs or BNLs) the corresponding tips belong to and the placement of RefPlantNLRs (Kourelis *et al*. 2021). Branch scale represents the number of substitutions per site.

**Figure S4.**
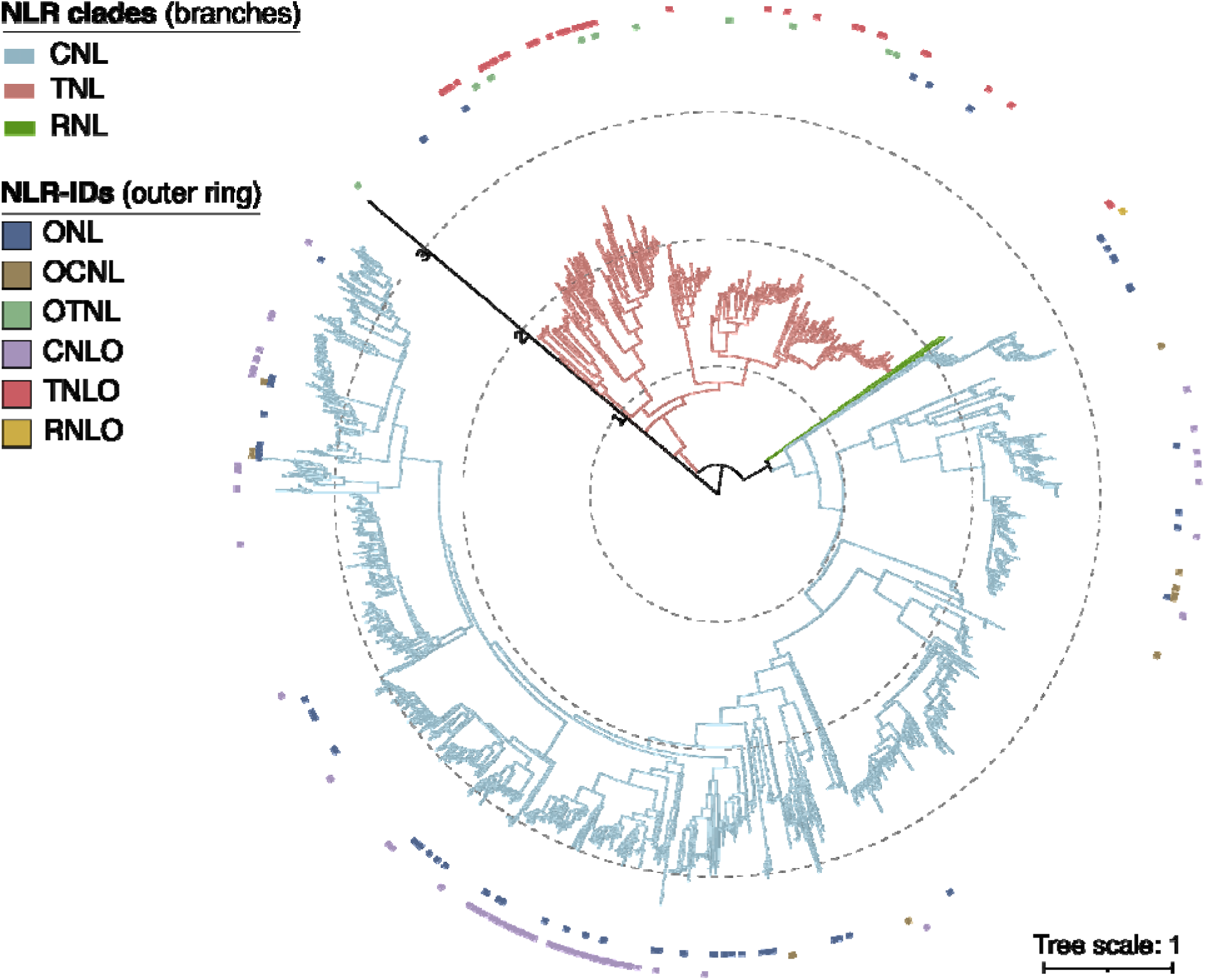
Sweetpotato and wild relatives harbor diverse sets of NLR-IDs clustering with canonical NLRs in the large NLR phylogeny. Maximum likelihood phylogeny of sweetpotato and wild relative NLRs inferred from central NB-ARC domain. Outer ring represents different types of NLR-IDs annotated by NLRtracker and their placement corresponds to their pair branch and model. The phylogeny was built using the Jones–Taylor–Thornton (JTT) and Per Site Rate (PSR) models in ExaML. The tree branches are rooted on the branch connecting TNL and non-TNL clades. The major TNL, CNL, and RNL clades are indicated by branch colors. Domain architecture and abbreviations are as shown in Figure S1. Branch scale represents the number of substitutions per site.

To examine the phylogenetic structure network of sensor and helper NLRs across the 35 genotypes, we extracted the CNL clade and examined the placement of well characterized NRCs including sensor and helper types. We highlighted the putative position of NRC0 and NRC1, which fall within a smaller clade in CNLs composed of 956 receptors (Figure 4). We defined this as the NRC-H subclade in sweetpotato and wild relatives, and extracted NLRs for further phylogenetic analysis. NRC0 clustered with 246 NLRs from sweetpotato and wild relatives. The majority of helper NLRs in our genotypes clustered separately (Figure 4). We labeled clades with known NRC sensors (NRC-S) and observed a radiating clade that branches from known NRC-S references like Bs2, Rx2, and Rpi-amr3i (Figure 4). Our phylogenetic analysis revealed a compact NRC-H subclade and an expanding NRC-S clade among sweetpotato genotypes and wild relatives.

**Figure 4.**
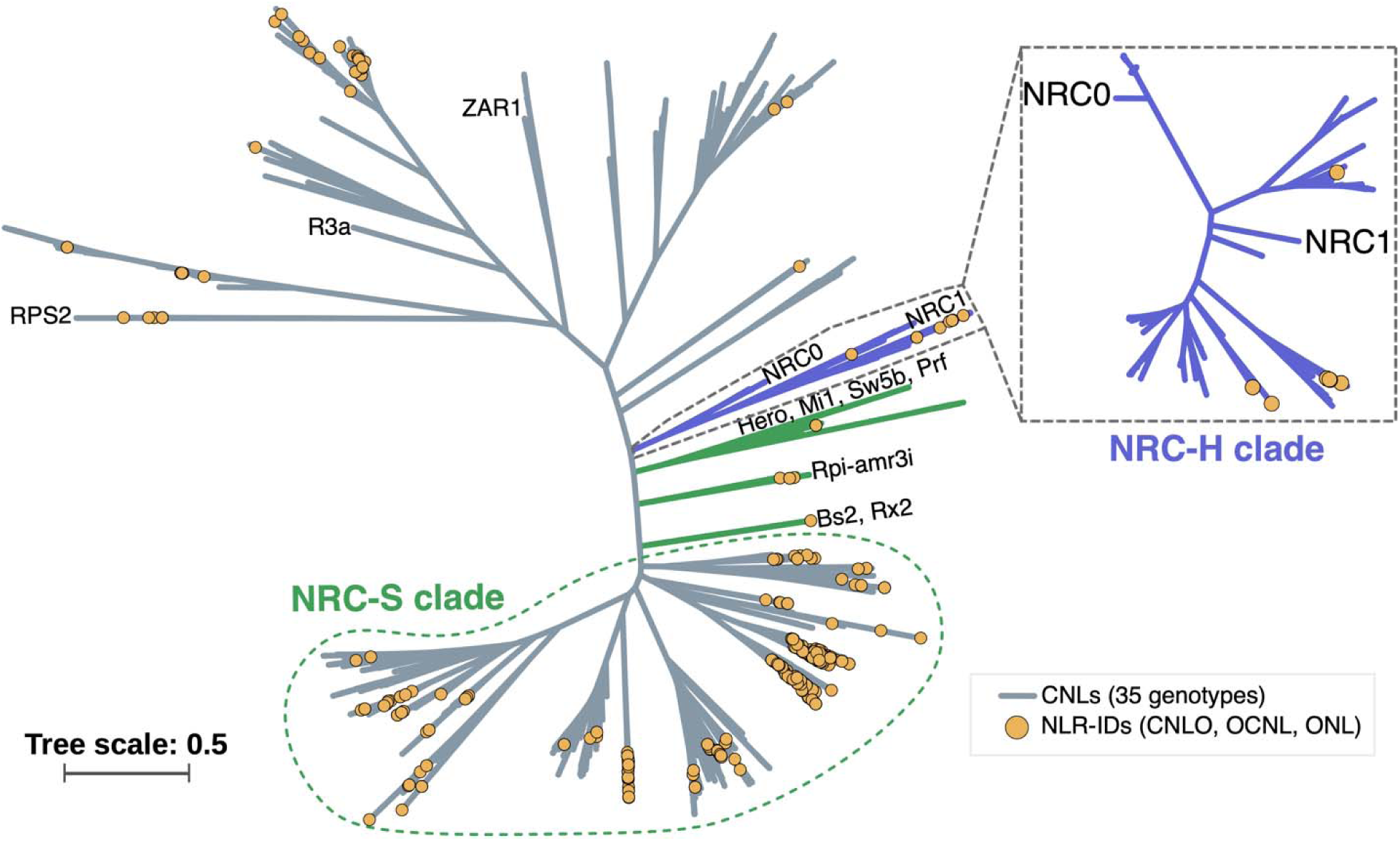
Sweetpotato genotypes and wild relatives harbor a compact NRC-H subclade. Phylogeny of sweetpotato and wild relative CNLs. The Maximum Likelihood tree includes only NB-ARC sequences corresponding to complete CNLs as predicted by NLRtracker. Branches predicted to correspond to major NLR required for cell death – helper and sensor clades (NRC-H, NRC-S) were highlighted based on phylogenetic placement of NRC0/1 (helpers-purple) and Hero-A, Rpi-amr3i, and Bs2 (sensors-green), respectively. Tips linked to CNLs containing integrated domains are labeled with yellow dots. The purple phylogenetic tree (right) includes only sequences from the indicated NRC-H lineage (left), underlining the *I. batatas*, *I. trifida*, *I. triloba* and *I. littoralis* sequences phylogenetically predicted as helper NLRs. Domain architecture and abbreviations are as shown in Figure S1.

### A substantial core of NLRs in sweetpotato

To understand the NLR allelic variation among sweetpotato genotypes, we clustered NLRs from all 32 sweetpotato genotypes into orthogroups (OGs) based on NB-ARC sequence identity at an optimal cutoff of 1.5% amino acid divergence (98.5% identity). We curated the sweetpotato NLRome into 4,366 OGs that corresponded to complete NLR domain architectures. We observed a cohort of singleton NLRs with 37 % of all OGs (N=1,626) falling into any single genotype, however singletons account for only 5.9% of the total NLR counts (N=27,615) across all sweetpotato genotypes. Figure 5 shows the classification of OGs based on the number of OGs among shared sweetpotato genotypes, their proportion across NLR counts, and their corresponding domain architecture. The core NLRome included 329 OGs (7.5%) shared by 21 to 32 genotypes. A cohort of 602 OGs (13.8%) were categorized as shared between 11 to 20 genotypes; we defined this cohort as the shell. The cloud constituted the largest group with 3,435 OGs (78.7%) found in 10 or fewer genotypes (Figure 5). We recorded 2,661 OGs consisting of CNLs, which corroborates with the highest abundance found by NLRtracker. A total of 50 OGs were shared by all 32 genotypes accounting for 1,600 NLRs (Figure 5). When examining the proportion of NLR counts within each of the categories and ignoring singletons, the core, shell, and cloud accounted for 33.84%, 33.47%, and 32.69% of the total NLR counts, respectively (Figure 5). Together, this analysis demonstrates the large conservation of NLRs in sweetpotato and corroborates that NLRs in sweetpotato present limited divergence at the individual genotype level.

**Figure 5.**
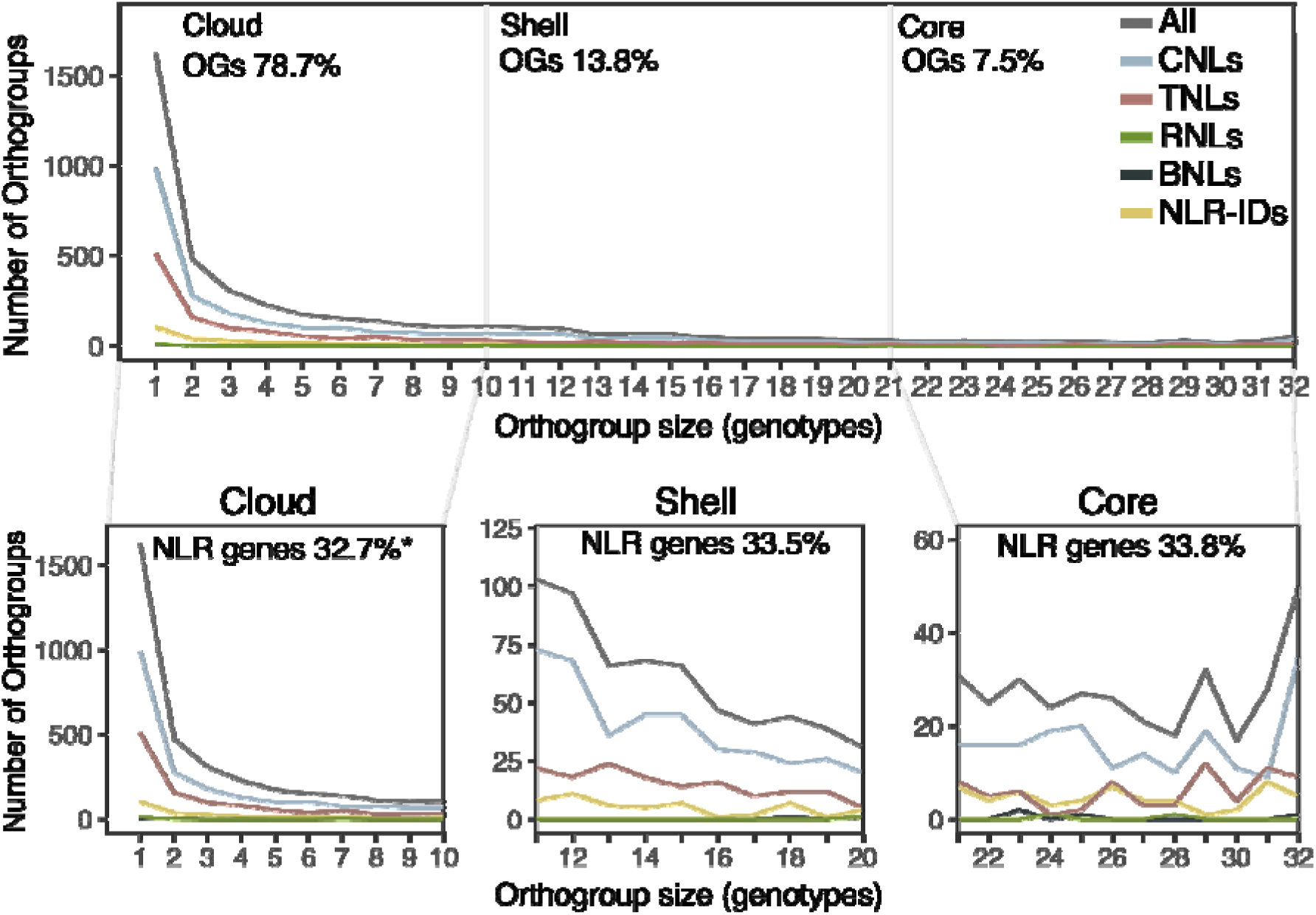
NLRs in sweetpotato present high conservancy. Orthogroup (OG) size distribution among 32 sweetpotato genotypes. Top line graph indicates the distribution of OGs shared by any of the 32 sweetpotato genotypes. NLR domain types associated with corresponding OGs are denoted by line colors. Percentage of OGs shared by genotypes in each category, Cloud (1 – 10 genotypes), Shell (10 – 20 genotypes) and Core (21 – 32 genotypes) are shown on top. Bottom line graphs show OG category specific distribution of sweetpotato NLRs in cloud (left), shell (center), and core (right). Percentage of NLR genes corresponding to each category is indicated on top. Asterisk (*) denotes that the NLR gene percentage was calculated excluding singletons. Domain architecture and abbreviations are as shown in Figure S1.

### CNLs among sweetpotato genotypes remain largely conserved

We documented large phylogenetic diversity and an expansion in the CNL clade among *Ipomoea* genotypes (Figure 3). To evaluate CNL conservation across sweetpotato genotypes, we used a phylogenetic tree from 32 sweetpotato genotypes and each of the 3 diploid wild relatives. We calculated the phylogenetic (patristic) distance between each of the 197 CNLs from *I. trifida*, 106 CNLs from *I. triloba*, 74 CNLs from *I. littoralis* to their closest phylogenetic neighbor from each of the 32 sweetpotato genotypes. We found that most CNLs in the 32 sweetpotato genotypes have short phylogenetic distance to their orthologs in the wild relatives (Figure 6, S5, and S6). This analysis revealed a set of 20 CNLs with patristic distance greater than 0.5 when compared against *I. trifida*, the progenitor of cultivated sweetpotato (Figure 6). A total of 13 CNLs from *I. triloba* and 8 CNLs for *I. littoralis* displayed greater than 0.5 phylogenetic distance when compared against all 32 sweetpotato genotypes (Figure S5 and S6). We identified a cluster of 4 CNLs that consistently exhibit high phylogenetic distance across all 32 sweetpotato genotypes against CNLs in *I. trifida*. Notably, a single CNL ortholog across Bwanjule, Murasaki-29, MC14-0363, NC07-0847, Porto Rico, Tib 11, and White Bonita exhibited the highest phylogenetic distance from its corresponding ortholog in *I. trifida* (Figure 6). Altogether, this analysis revealed that a small repertoire of *I. batatas* CNLs diverged from *I. trifida*, *I. triloba*, and *I littoralis*.

**Figure 6.**
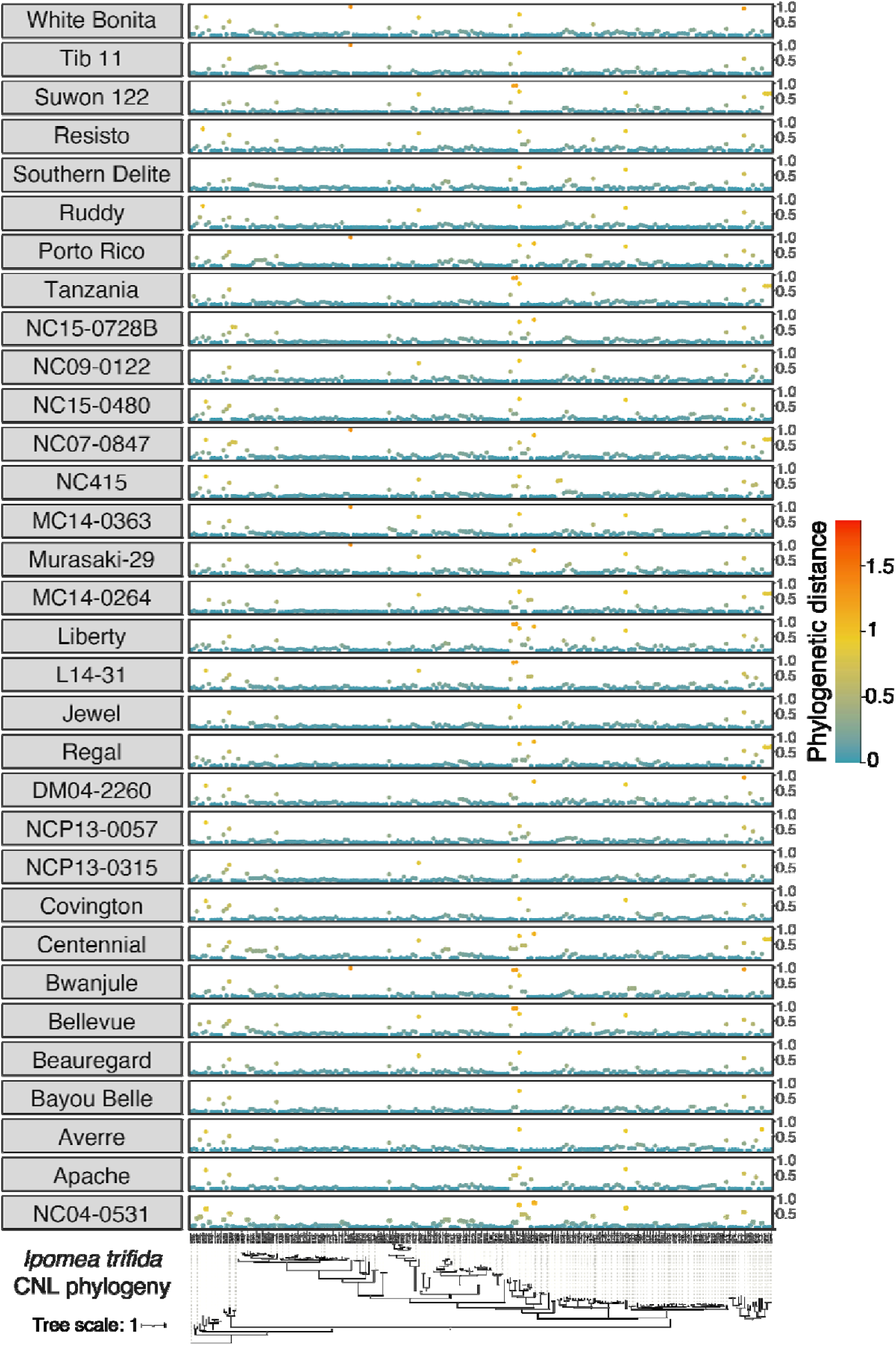
Widespread CNL conservation between *I. trifida* and *I. batatas* genotypes (N= 32) Phylogenetic (patristic) distance of two CNL nodes between *I. trifida* and each *I. batatas* genotypes were calculated from a combined NLR phylogeny. The patristic distances for each corresponding *I. batatas* CNLs are plotted with color scale indicating the distance level for each pair. Domain architecture and abbreviations are as shown in Figure S1.

**Figure S5.**
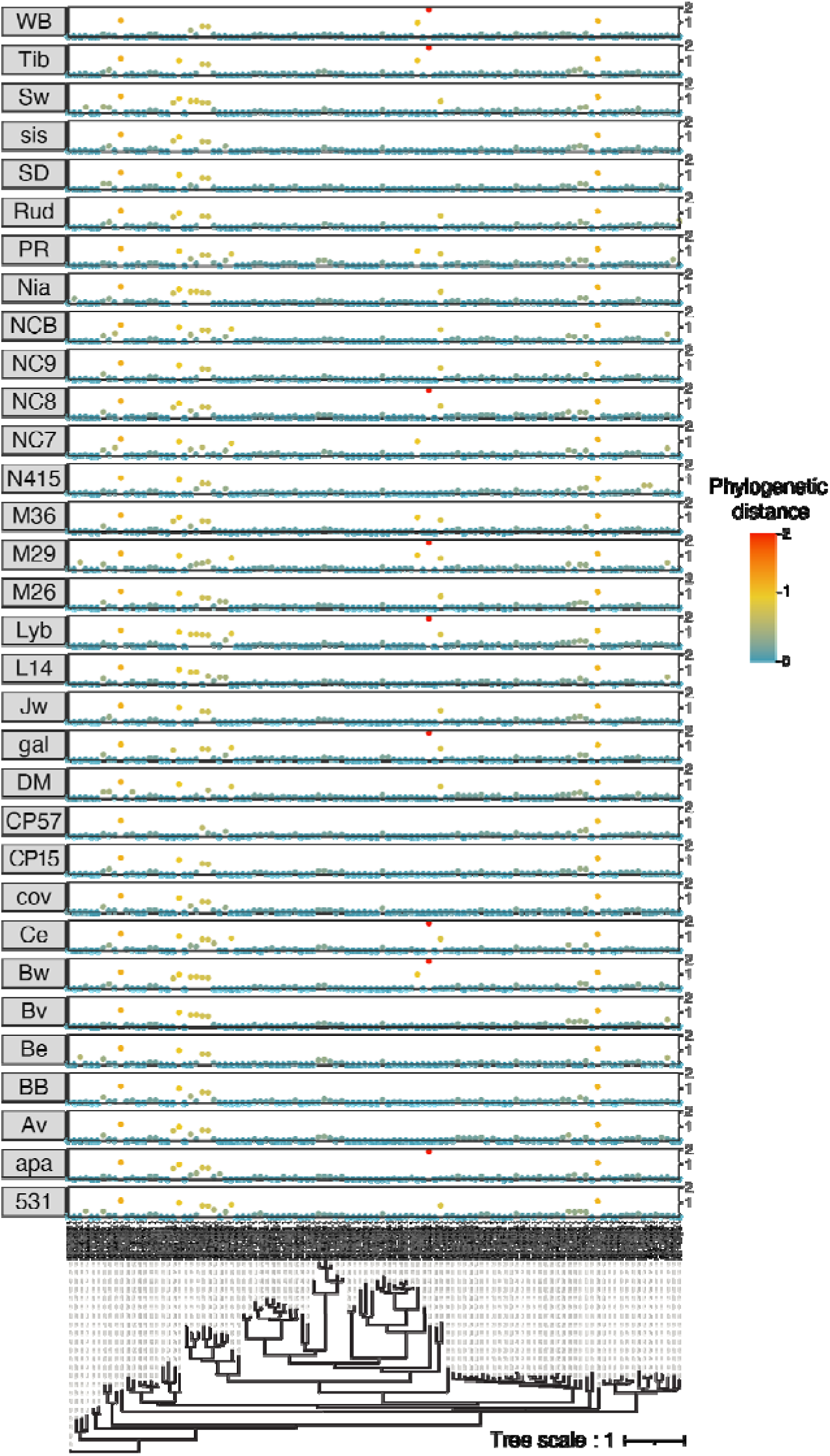
CNL conservation between *Ipomoea triloba* and *I. batatas* genotypes (N= 32) Phylogenetic (patristic) distance of two CNL nodes between *I. triloba* and each *I. batatas* genotypes were calculated from a combined NLR phylogeny. The patristic distances for each corresponding *I. batatas* CNLs are plotted with color scale indicating the distance level for each pair. Genotype abbreviations match names contained in Table S4. Domain architecture and abbreviations are as shown in Figure S1.

**Figure S6.**
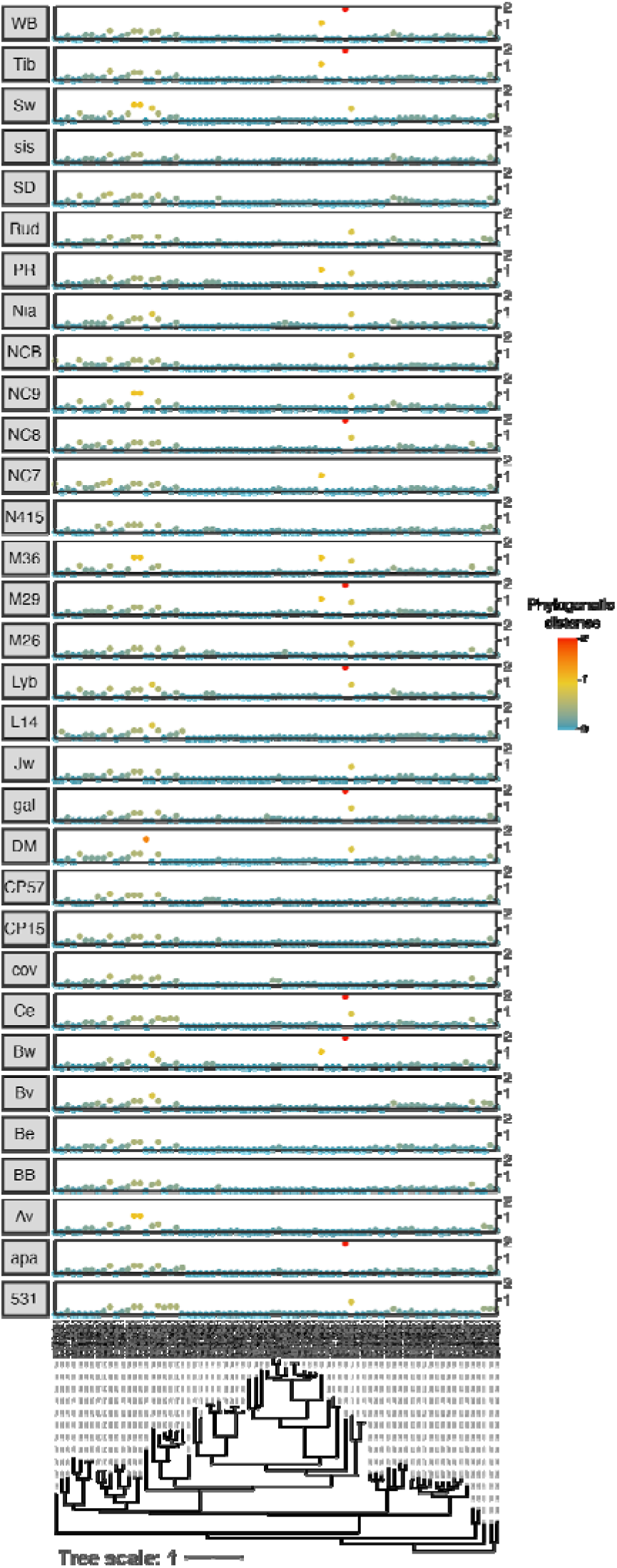
CNL conservation between *Ipomoea littoralis* and *I. batatas* genotypes (N= 32) Phylogenetic (patristic) distance of two CNL nodes between *I. littoralis* and each *I. batatas* genotypes were calculated from a combined NLR phylogeny. The patristic distances for each corresponding *I. batatas* CNLs are plotted with color scale indicating the distance level for each pair. Genotype abbreviations match names contained in Table S4. Domain architecture and abbreviations are as shown in Figure S1.

### Anchoring NLR loci in chromosome-level genomes of I. batatas and I. trifida

To pinpoint areas in the genome harboring NLRs, we positioned Beauregard and *I. trifida* NLR loci on the 90 and 15 chromosome scale genome assemblies for *I. batatas* and *I. trifida*, respectively (Figure 7 and 8). With a threshold of 99% identity and focusing only on NLRs contigs associated to NLRs with complete NLR architecture, we placed 810 NLR contigs for *I. batatas* and 313 contigs for *I. trifida*. These contigs represent major NLR domain architectures. We recorded clustering of CNLs in *I. batatas* Chromosomes 4, 9, 10, 11, 13, and 14 and *I. trifida*’s Chromosomes 1, 3, 4, 9, 10, 11, 13, 14, and 15 (Figure 7 and 8). Contigs associated with TNLs clustered in Chromosome 7 and 12 for both *I. batatas* and *I. trifida*. We observed remarkable clustering at the distal portion of chromosomes. For *I. batatas*, some chromosome copies completely lacked NLR loci including Chromosome 1D, 2B, 5C, 5A, 6A, 6F, 8C, 8D, 8E, 11D, 12F, and 13B. Together, these results represent the state-of-the-art physical mapping of NLR loci on the hexaploid Beauregard *I. batatas* and the diploid *I. trifida* assembly.

**Figure 7.**
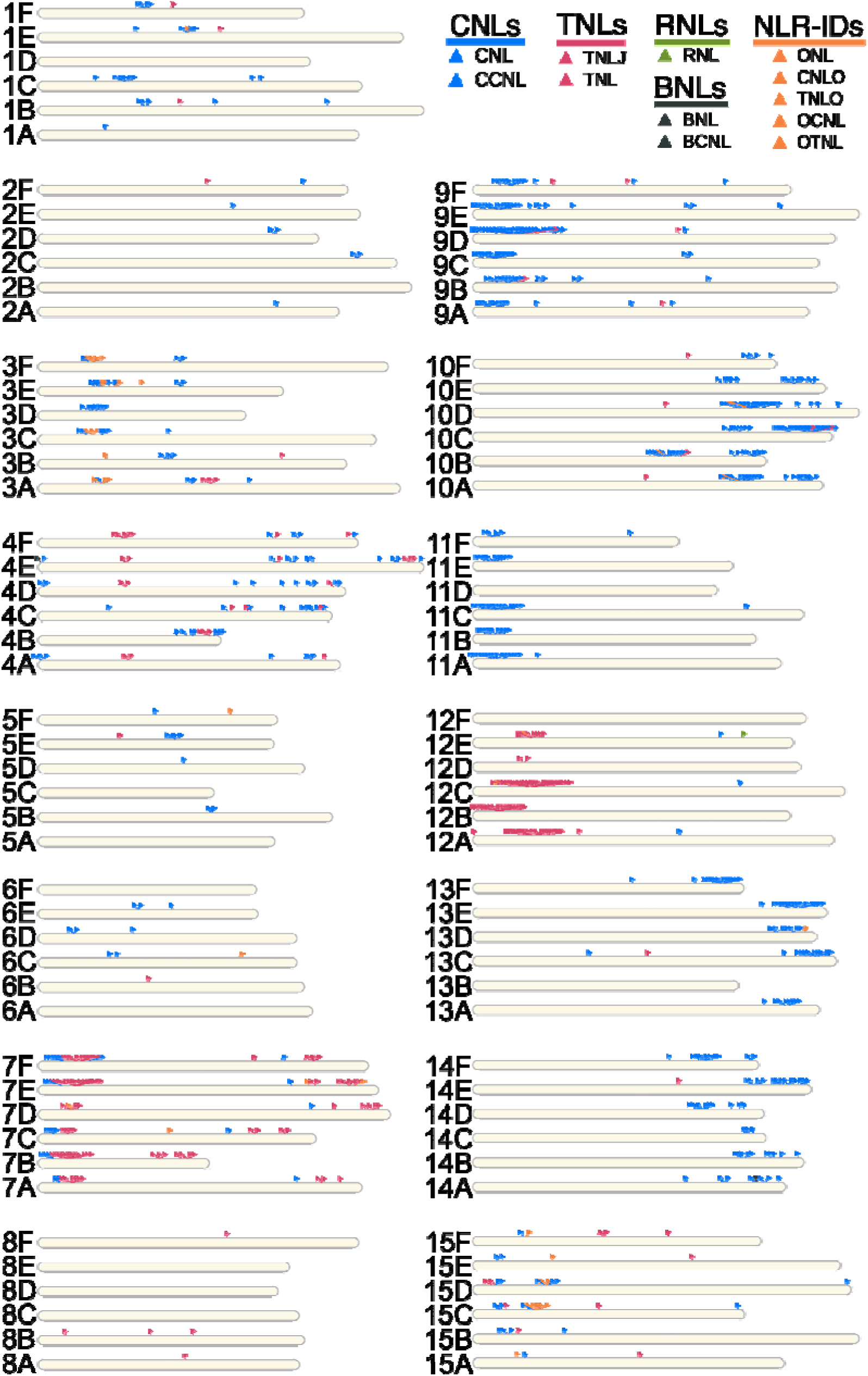
RenSeq allowed anchoring of NLR contigs corresponding to complete NLR domain architecture in *Ipomoea batatas*. Physical positions of sweetpotato genotype Beauregard NLR contigs displayed along the 90 chromosome diagrams for the Beauregard genome assembly. Each contig is represented by a triangle marked with colors corresponding to the associated NLR architectures. Blue triangles correspond to CNLs, coiled-coil nucleotide-binding and leucine-rich repeat immune receptors (i.e. CNL or CCNL); dark grey triangles correspond to BNLs, Late-Blight R1 nucleotide-binding and leucine-rich repeat immune receptors (i.e. BNL or BCNL); red triangles are for TNLs, Toll/interleukin-1 receptor nucleotide-binding and leucine-rich repeat immune receptors with or without C-terminal jelly roll/Ig-like domain (i.e. TNL or TNLJ); green triangles correspond to RNLs, N-terminal RPW8-type coiled-coil nucleotide-binding and leucine-rich repeat immune receptors; and orange triangles correspond to NLR-IDs, nucleotide-binding and leucine-rich repeat immune receptors containing non canonical “integrated domains”. Detailed domain architecture and abbreviations are as shown in Figure S1.

**Figure 8.**
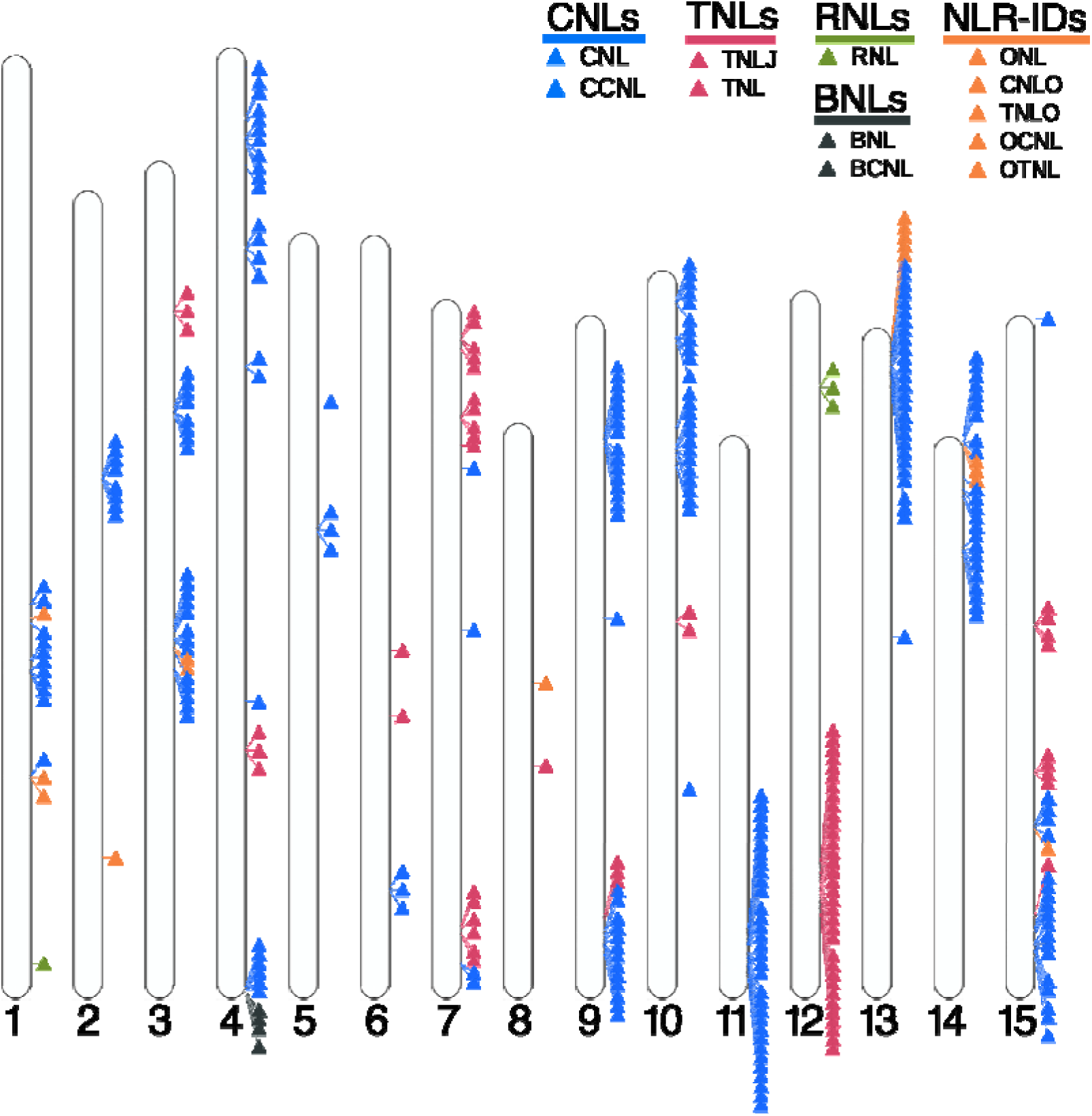
RenSeq allowed anchoring of NLR contigs corresponding to complete NLR domain architecture in *Ipomoea trifida*. Physical positions of *I. trifida* NLR contigs displayed along the 15 chromosome diagrams for the *I. trifida* genome assembly. Each contig is represented by a triangle marked with colors corresponding to the associated NLR architecture. Blue triangles correspond to CNLs, coiled-coil nucleotide-binding and leucine-rich repeat immune receptors (i.e. CNL or CCNL); dark grey triangles correspond to BNLs, Late-Blight R1 nucleotide-binding and leucine-rich repeat immune receptors (i.e. BNL or BCNL); red triangles are for TNLs, Toll/interleukin-1 receptor nucleotide-binding and leucine-rich repeat immune receptors with or without C-terminal jelly roll/Ig-like domain (i.e. TNL or TNLJ); green triangles correspond to RNLs, N-terminal RPW8-type coiled-coil nucleotide-binding and leucine-rich repeat immune receptors; and orange triangles correspond to NLR-IDs, nucleotide-binding and leucine-rich repeat immune receptors containing non canonical “integrated domains”. Detailed domain architecture and abbreviations are as shown in Figure S1.

## MATERIALS AND METHODS

### Plant material, growth conditions and DNA extractions

To capture the global and local diversity of *Ipomoea batatas,* we included 32 hexaploid *I. batatas* genotypes and three diploid wild *Ipomoea* sp. genotypes. We selected a set of 32 representative *I. batatas* genotypes based on their importance and potential as parents of mapping populations. This *I. batatas* panel included land races, cultivated, and advanced breeding lines. We also included three wild *Ipomoea* species including *I. littoralis* (PI 573335), *I. triloba* (NCNSP0323), and *I. trifida* (NCNSP0306); the former is considered the progenitor of cultivated sweetpotato (Table S5 and Methods S1). Genomic DNA of young leaf tissues obtained from one to two plants per genotype was extracted using NucleoBond HMW DNA Kit (MACHEREY - NAGEL Inc., PA, USA). We extracted approximately 10 µg of genomic DNA per genotype to allow downstream library preparation.

**Table S5.**
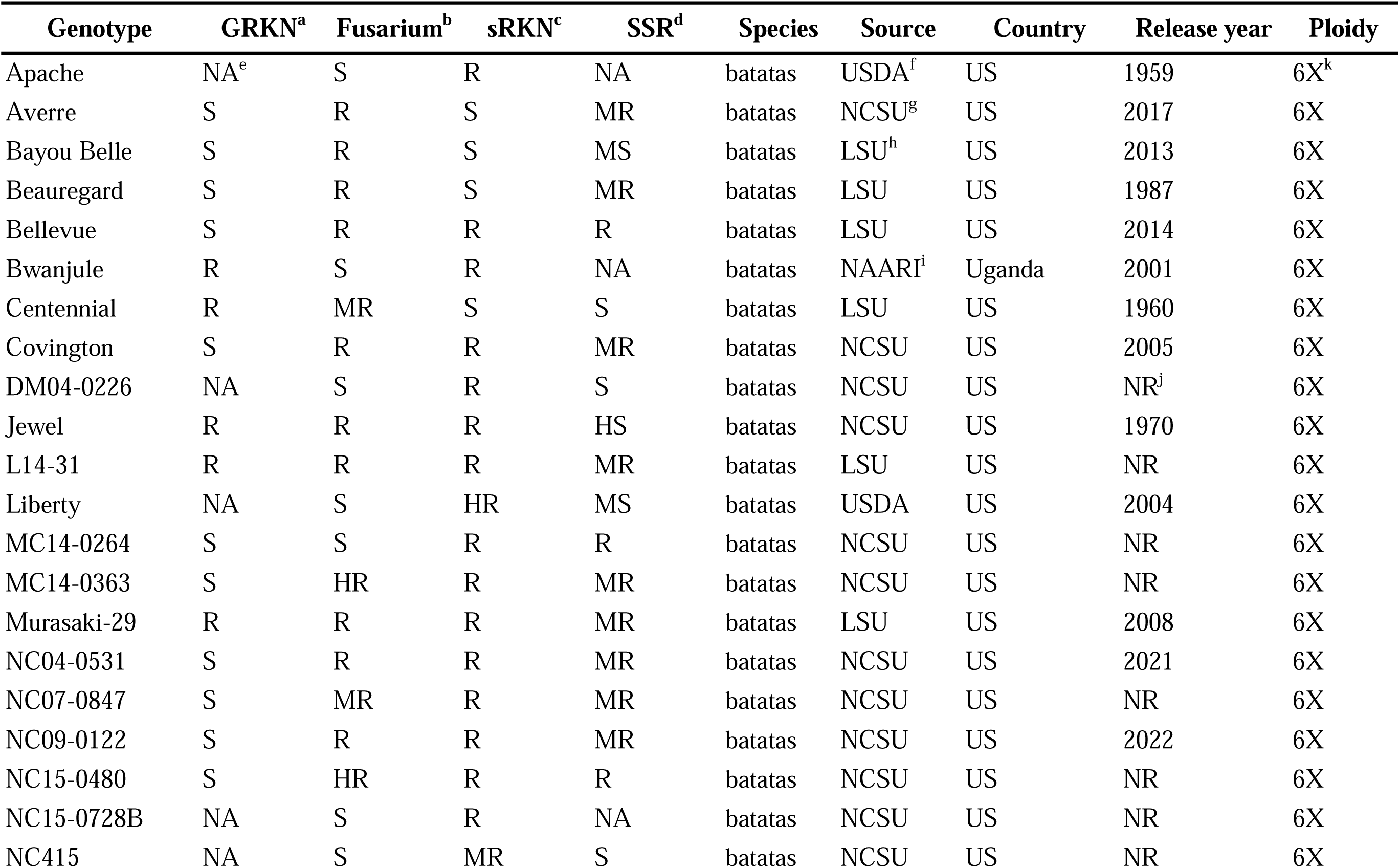
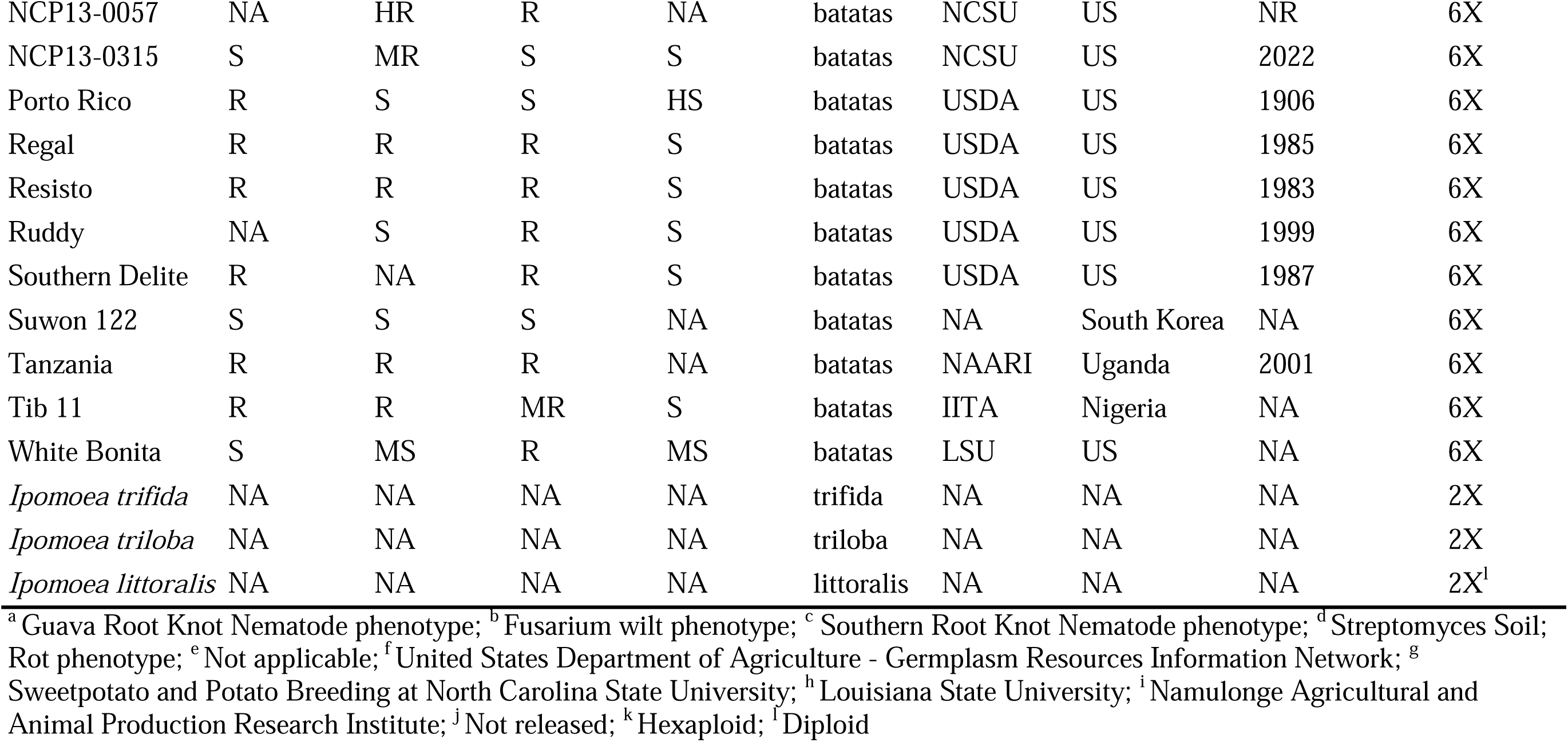
Metadata information for 32 sweetpotato genotypes and 3 wild relatives included in our RenSeq experiment.

| Genotype | GRKN<sup>a</sup> | Fusarium<sup>b</sup> | sRKN<sup>c</sup> | SSR<sup>d</sup> | Species | Source | Country | Release year | Ploidy |
| --- | --- | --- | --- | --- | --- | --- | --- | --- | --- |
| Apache | NA<sup>e</sup> | S | R | NA | batatas | USDA<sup>f</sup> | US | 1959 | 6X<sup>k</sup> |
| Averre | S | R | S | MR | batatas | NCSU<sup>g</sup> | US | 2017 | 6X |
| Bayou Belle | S | R | S | MS | batatas | LSU<sup>h</sup> | US | 2013 | 6X |
| Beauregard | S | R | S | MR | batatas | LSU | US | 1987 | 6X |
| Bellevue | S | R | R | R | batatas | LSU | US | 2014 | 6X |
| Bwanjule | R | S | R | NA | batatas | NAARI<sup>i</sup> | Uganda | 2001 | 6X |
| Centennial | R | MR | S | S | batatas | LSU | US | 1960 | 6X |
| Covington | S | R | R | MR | batatas | NCSU | US | 2005 | 6X |
| DM04-0226 | NA | S | R | S | batatas | NCSU | US | NR<sup>j</sup> | 6X |
| Jewel | R | R | R | HS | batatas | NCSU | US | 1970 | 6X |
| L14-31 | R | R | R | MR | batatas | LSU | US | NR | 6X |
| Liberty | NA | S | HR | MS | batatas | USDA | US | 2004 | 6X |
| MC14-0264 | S | S | R | R | batatas | NCSU | US | NR | 6X |
| MC14-0363 | S | HR | R | MR | batatas | NCSU | US | NR | 6X |
| Murasaki-29 | R | R | R | MR | batatas | LSU | US | 2008 | 6X |
| NC04-0531 | S | R | R | MR | batatas | NCSU | US | 2021 | 6X |
| NC07-0847 | S | MR | R | MR | batatas | NCSU | US | NR | 6X |
| NC09-0122 | S | R | R | MR | batatas | NCSU | US | 2022 | 6X |
| NC15-0480 | S | HR | R | R | batatas | NCSU | US | NR | 6X |
| NC15-0728B | NA | S | R | NA | batatas | NCSU | US | NR | 6X |
| NC415 | NA | S | MR | S | batatas | NCSU | US | NR | 6X |
| NCP13-0057 | NA | HR | R | NA | batatas | NCSU | US | NR | 6X |
| NCP13-0315 | S | MR | S | S | batatas | NCSU | US | 2022 | 6X |
| Porto Rico | R | S | S | HS | batatas | USDA | US | 1906 | 6X |
| Regal | R | R | R | S | batatas | USDA | US | 1985 | 6X |
| Resisto | R | R | R | S | batatas | USDA | US | 1983 | 6X |
| Ruddy | NA | S | R | S | batatas | USDA | US | 1999 | 6X |
| Southern Delite | R | NA | R | S | batatas | USDA | US | 1987 | 6X |
| Suwon 122 | S | S | S | NA | batatas | NA | South Korea | NA | 6X |
| Tanzania | R | R | R | NA | batatas | NAARI | Uganda | 2001 | 6X |
| Tib 11 | R | R | MR | S | batatas | IITA | Nigeria | NA | 6X |
| White Bonita | S | MS | R | MS | batatas | LSU | US | NA | 6X |
| *Ipomoea trifida* | NA | NA | NA | NA | trifida | NA | NA | NA | 2X |
| *Ipomoea triloba* | NA | NA | NA | NA | triloba | NA | NA | NA | 2X |
| *Ipomoea littoralis* | NA | NA | NA | NA | littoralis | NA | NA | NA | 2X<sup>1</sup> |

<sup>a</sup> Guava Root Knot Nematode phenotype; <sup>b</sup> Fusarium wilt phenotype; <sup>c</sup> Southern Root Knot Nematode phenotype; <sup>d</sup> Streptomyces Soil; Rot phenotype; <sup>e</sup> Not applicable; <sup>f</sup> United States Department of Agriculture - Germplasm Resources Information Network; <sup>g</sup> Sweetpotato and Potato Breeding at North Carolina State University; <sup>h</sup> Louisiana State University; <sup>i</sup> Namulonge Agricultural and Animal Production Research Institute; <sup>j</sup> Not released; <sup>k</sup> Hexaploid; <sup>l</sup> Diploid

### NLR gene enrichment sequencing

To design our target NLR bait library, we used NLR-parser, a benchmarked NLR annotation tool, that provides sequence coordinates of complete and partial NLRs in a set of query sequences (Steuernagel *et al*. 2015). We scanned the available genomic resources for sweetpotato, including two diploid wild relatives (*I. trifida* and *I. triloba*) high-confidence coding DNA sequences (cDNA), a transcriptome assembly from the hexaploid genotype Beauregard, and cDNA from *I. nil* (Steuernagel *et al*. 2015). The cDNA sequences represent spliced transcript models, including untranslated regions, and were chosen to ensure comprehensive representation of gene models. Only complete putative NLR sequences were used to create a bait library as described by Jupe *et al*. (2013). In brief, the library was composed of 38,694 120-mer biotinylated RNA baits starting from the first nucleotide following the predicted coding region and synthesized by Arbor Biosciences (Arbor Biosciences, Ann Arbor, MI, USA). A total of 10 µg of high molecular weight (HMW) genomic DNA (gDNA) from each genotype were fragmented with a sonicator to obtain 3-5 Kb fragments (Covaris Inc., Woburn, MA, USA). NLR capture followed MYbaits v4.0 protocol with modifications detailed in Methods S1. To test the enrichment efficiency, we performed qPCRs on four NLR targets included in the bait library and Actin gene. Quantitative PCRs (qPCRs) primers are listed in Table S6 and protocols are detailed in Methods S1. Captured libraries that passed the enrichment efficiency check (8 - 10 cycle difference) were amplified using high fidelity KAPA enzyme 1 U/µL (Roche, Indianapolis, IN) and subsequently prepared for PacBio SMRT sequencing following the NC State-GSL (Genomic Sciences Laboratory) standard recommendations for 4-10 kb library preparation. The libraries were sequenced using the Sequel PacBio platform at the NC State GSL.

**Table S6.** Primers used for enrichment efficiency check qPCR. *https://doi.org/10.6084/m9.figshare.21970805*

### Assembly, structural and functional NLR annotation

Circular Consensus Sequencing (CCS) reads were generated from raw subreads using *ccs* with three full passes and 90% accuracy (PacBio, 2022). Each set of CCS reads from each genotype was processed to remove the adapters and barcodes using Cutadapt (version 1.16) (Martin, 2011). To assess the number of CCS reads containing at least one NLR bait sequence, we conducted a BLAST search of the entire NLR bait library on each genotype CCS read library using BLAST+ (v.2.9.0). We also calculated the number of reads containing NLR motifs using NLR-parser to scan DNA sequences (Steuernagel *et al*. 2015). Only non-chimeric reads were assembled using Canu (version 1.6) (Koren *et al*. 2017). We reported the number of contigs containing complete NLR motifs as defined by NLR-annotator (version 1.0) (Zhang, 2020). All contigs were retained during annotation to preserve haplotypic variants and ensure the accurate representation of allelic diversity. A custom MAKER2 annotation pipeline was designed to predict NLR gene models (version 2.31.9) (Holt and Yandell, 2011). Protein evidence from *I. trifida, I. triloba* and *I. nil* was externally aligned using Exonerate (version 2.2.0). Transcript evidence consisted of Beauregard ONT (Oxford Nanopore Technology) full-length cDNA reads generated from combined leaf, fibrous, and storage root tissues (Buell *unpublished*). Gene predictions from representative hexaploid (Beauregard) and diploid (*I. trifida)* were used to train AUGUSTUS (Korf, 2004) and SNAP HMMs (Hidden Markov Models) (Stanke *et al*. 2008). Evidence and gene model inspection was carried out in the Integrative Genomics Viewer software (IGV)(version IGV 2.14.x) (Robinson *et al*. 2011). NLR protein models were classified based on their multi-domain architecture using the benchmarked NLRtracker tool (Kourelis *et al*. 2021). We arbitrarily set a threshold to select complete NLR protein models that consisted of: models carrying the canonical NB-ARC, LRR, and one the N or C-terminal domains (CC, TIR, RPW8, B, CID-J, and IDs) (Figure **S1**). The current notion of integrated domains suggest that NLRs carrying IDs represent key pathogen effector targets, therefore we looked at their distribution in our dataset by grouping them into 6 distinct subclasses: ONL, OCNL, CNLO, OTNL, TNLO, and RNLO (Figure **S1**) (Cesari *et al*. 2014). Greater detail is provided in Methods S1. Table S4 lists the general abbreviation/naming convention for each of the 35 genotypes included in this study.

**Figure S1.**
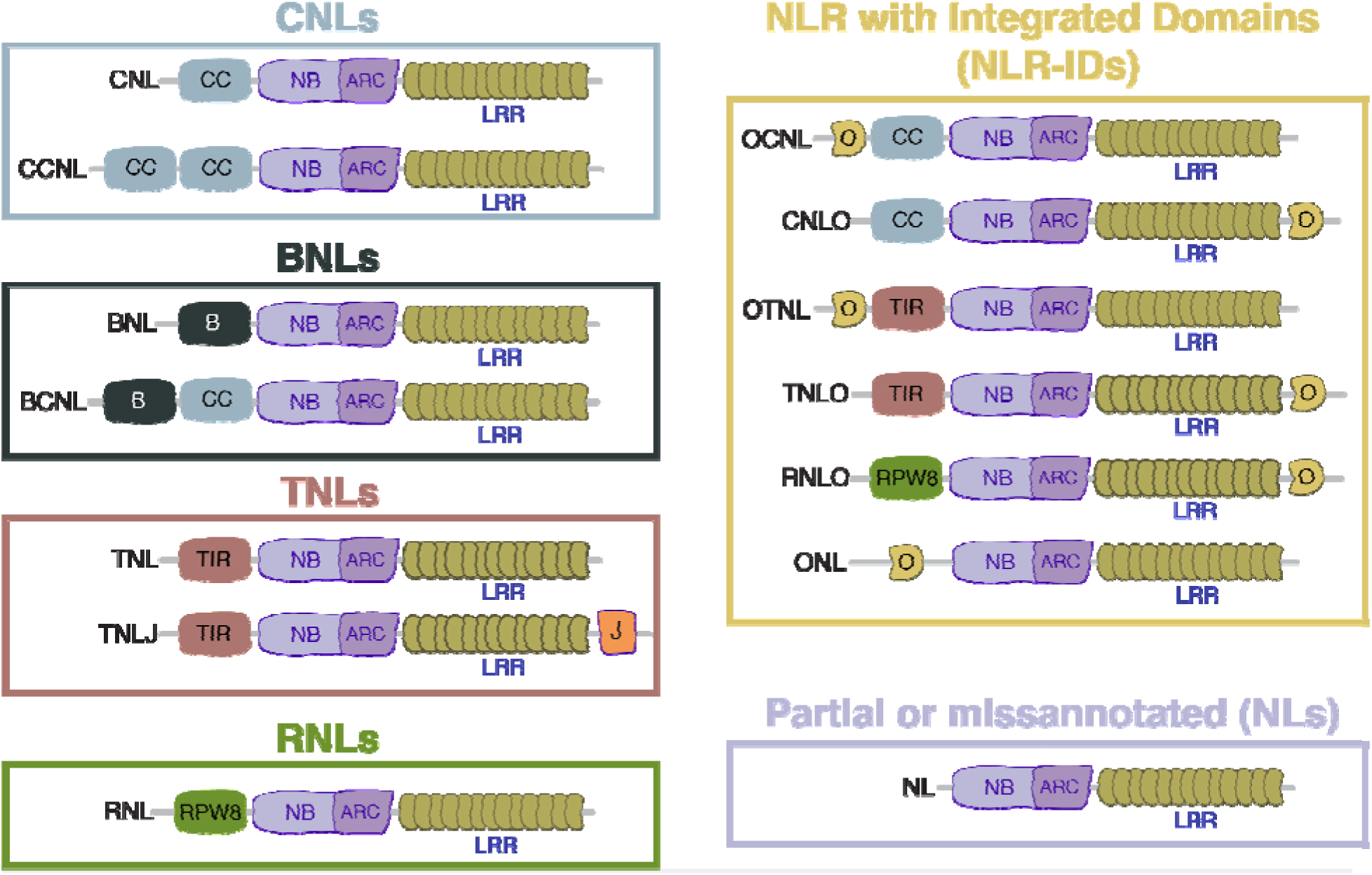
Modular representation of NLR domain architecture diversity examined in this study. NB-ARC, nucleotide-binding adaptor shared by APAF-1; CNLs, coiled-coil nucleotide-binding and leucine-rich repeat immune receptors (i.e. CNL or CCNL); BNLs, Late-Blight R1 nucleotide-binding and leucine-rich repeat immune receptors (i.e. BNL or BCNL); TNLs, Toll/interleukin-1 receptor nucleotide-binding and leucine-rich repeat immune receptors with or without C-terminal jelly roll/Ig-like domain (i.e. TNL or TNLJ); RNLs, N-terminal RPW8-type coiled-coil nucleotide-binding and leucine-rich repeat immune receptors; NLR-IDs, nucleotide-binding and leucine-rich repeat immune receptors containing non canonical “integrated domains”. The integrated domain may reside at the N- or C-terminus of the protein. We defined partial or missannotated NLRs as those models comprise of NB-ARC and LRR domains (NLs).

**Table S4.** Gene model naming scheme. This table shows our abbreviation scheme and our fasta header gene ID scheme. Notice that for example a gene id “iba_apa00730g01.1” includes the following information: iba = Ipomoea batatas; apa= genotype apache; 00730 = contig number; g01= specifies the gene number in the respective contig ex. g01 means this is the first gene in the contig; .1 = remind us that this gene has a transcript sequence associated in a different file. *https://doi.org/10.6084/m9.figshare.21899877*

### Comparison of NLR content in standard genome annotation

To highlight the differences in NLR annotation outcomes between standard genome annotation projects and RenSeq/NLR tailored annotation pipelines, the proteomes from 8 plant species were downloaded from individual genome project repositories (Methods S1). All protein sequences were scanned through NLRtracker and categorized as NLRs if the sequence contained an NB-ARC domain and at least 1 additional domain. We compared each genome annotation NLR count with that reported for the corresponding RenSeq projects for the same plant species. The NLR counts from RenSeq projects were collected as reported in each RenSeq study.

### NLR phylogenetic diversity in 6X and 2X *Ipomoea* spp

To explore the diversity of NLRs in the 32 *I. batatas* hexaploid and 3 diploid wild relatives, we constructed NLR phylogenies for complete NLRs only. We employed NB-ARC domains extracted by NLRtracker to produce a multiple sequence alignment (MSA). We included 35 refPlantNLRs that encompasses major functionally annotated and phylogenetically diverse NLRs (Kourelis et al. 2021). The MSA was generated using the globalpair alignment in MAFFT (version 7.490) (Rozewicki *et al*. 2019). A maximum-likelihood tree was inferred from the resulting MSA of 379 columns and 29,553 sequences using ExaML (version 3.0.17) (Kozlov *et al*. 2015). We inferred 6 randomized stepwise addition order parsimony-based starting trees required for ExaML using RAxML (version 8.1.20) (Stamatakis, 2014). The final tree was visualized in the Interactive Tree Of Life (iTOL) software (version 6.6) and arbitrarily rooted on the branch connecting TNLs and non-TNL clades (Letunic and Bork, 2007). We pruned and extracted NLRs that clustered with functionally characterized NRC0 and NRC1 NLRs and visualized the phylogeny in iTOL (Methods S1).

### Phylogenetic distance analysis

To evaluate NLR conservation between diploid wild relatives (N = 3) and *I. batatas* genotypes (N= 32), we calculated phylogenetic distance among complete CNLs as they represent the largest expanding NLR clade in our study. First, we extracted NB-ARC deduplicated domains corresponding to complete CNLs in all 35 genotypes. A total of 6 NB-ARC datasets were generated; three datasets represented the 3 wild relatives individually (*I. trifida*, *I. triloba*, and *I. littoralis*) and the other 3 datasets included all 32 hexaploid *I. batatas* genotypes in combination with a single wild relative (32 + Itf; 32 + Itb; 32 + lito). We aligned the NB-ARC amino acid sequences using MAFFT (version 7.505). We generated an NLR phylogeny for each dataset described above. We calculated the phylogenetic (patristic) distance between each pair of CNLs in the diploid wild relatives and their corresponding closest CNLs in the 32 hexaploid sweetpotato genotypes. We visualized distance against each diploid wild relative phylogeny in R using ggplot2 (Methods S1).

### Orthology inference, refinement, and classification

We inferred orthologous groups using a pairwise global amino acid similarity approach over the length of the NB-ARC domains extracted by NLRtracker from each of the 35 genotypes using BLAST+ (v.2.9.0) (Methods S1). For any pair of genes with a BLAST Evalue > 1e−20, we used the global alignment algorithm of Needleman and Wunsch (1970) to align the two sequences: we then computed the pairwise amino acid percent difference between the two sequences and stored this value in the matrix. We created a graph where edges connected nodes (sequences) with percent identity >98.5%. We then inferred orthogroups to be connected components within this graph. We kept orthogroups that contained sequences assigned to complete NLR domains (RNLO, TNLO, CNLO, OCNL, OTNL, ONL, BNL, BCNL, CNL, TNL, RNL, CCNL, and TNLJ) (Figure **S1**). The orthogroup counts per genotype were converted into a presence/absence matrix to examine orthogroup distribution among 32 sweetpotato genotypes and assign cloud, shell, and core categories. We classified orthogroups into the cloud category if the orthogroups were shared by < 10 genotypes, the shell category included orthogroups shared between 11 and 20 genotypes, finally the core orthogroup category included orthogroups shared by > 21 genotypes. The co-occurrence of orthogroups and their corresponding category across genotypes was visualized in R using ggplot2.

### Genomic anchoring of complete NLRs in hexaploid Beauregard

To examine the chromosome level clustering and location of NLRs in hexaploid sweetpotato, we anchored the hexaploid Beauregard RenSeq contigs into the recently haplotype-resolved chromosome-scale Beauregard genome assembly (*pre-publication version*) (Methods S1). We visualized NLR contigs along chromosomes using the R package RIdeogram (version 0.2.2) (Hao *et al*. 2020). We repeated this process for the *I. trifida* RenSeq contigs and its corresponding genome assembly (Wu *et al*. 2018b).

### Data availability

The CCS reads described here were deposited as raw data in the National Center for Biotechnology Information under the BioProject accession <u>PRJNA946648</u>. The metadata table linking SRA numbers, Biosample, and sweetpotato genotype information was deposited in fishare under the link (*https://doi.org/10.6084/m9.figshare.27872427*). All resulting NLRtracker annotations were deposited in FigShare (https://figshare.com/account/home#/projects/157356). All the assemblies were made available to the community in figshare (*https://doi.org/10.6084/m9.figshare.27635481*). We provided the FASTA file containing filtered and clustered NLR baits used in the sweetpotato RenSeq experiment (*https://doi.org/10.6084/m9.figshare.25303204*).

## DISCUSSION

In this study, we used RenSeq, a genome complexity reduction approach, to reveal a myriad of NLRs harbored in the genomes of 32 hexaploid sweetpotato genotypes and three diploid wild relatives. We captured, sequenced, and annotated a breadth of NLRs for all genotypes, with minimal off-target rate (2.7 %) to produce state-of-the-art NLR annotations. Early RenSeq projects obtained lower capture efficiency at the read and contig level which speaks to the quality of our bait design, skilled library preparation, and improvements by the bait library manufacturer (Giolai *et al*. 2016). When comparing genome projects versus SMRT RenSeq and NLR tailored annotations, we captured and annotated more NLRs than the genome annotation projects for *I. batatas*, *I trifida*, and *I .triloba* (Wu *et al*. 2018b). Our annotation pipeline utilized long read cDNA as evidence and avoided masking repetitive sequences during gene model predictions. Repeat masking introduces bias when annotating highly repetitive NLRs in Brassicaceae as documented by Bayer *et al*. (2018), who reported that *ab initio* annotation programs failed to distinguish transposable elements fused with NLR genes. Given the ploidy level (6X) and importance of this staple crop, our sweetpotato NLR repertoire exhibits great potential to advance sweetpotato resistance breeding and highlights the benefit of deploying RenSeq in other polyploid crops.

The CNL domain, which generally carries a coiled-coil motif at its N-terminus, ranked as the most common NLR domain in all wild and cultivated genotypes. In agreement with this observation, other Solanales species with high quality NLRomes also exhibit an expansion of CNLs (Jupe *et al*. 2013; Stam *et al*. 2016; Witek *et al*. 2016; Seong *et al*. 2020, 2022). Recently, Seong et al. (2020) reported the diversification of CNLs in 16 accessions from five different wild tomato relatives, in addition to *Nicotiana benthamiana* and *Capsicum annuum*, also belonging to the Solanales order. Our comparative phylogenetic analysis revealed separation of canonical domains as evidenced by our unrooted phylogeny and supported the diverging evolutionary history of the TNL and CNL clades as previously reported (Kourelis *et al*. 2021). Our NLR phylogeny confirmed the expansion of CNLs with a large number of tips radiating beyond any RefPlantNLRs included as reference. The majority of ONLs clustered within the expanding CNL clades and few within the TNL clade. Our sweetpotato NLR phylogeny allows for placement of several ONLs that would otherwise be unclassified.

The new paradigm of CNL networking categorizes NLRs into “sensor NLRs”, which function in direct recognition of pathogen effectors and “helper NLRs”, which interact with the sensors and mediate downstream immune signaling (Wu *et al*. 2018a; Adachi *et al*. 2019; Contreras *et al*. 2022). As helper and sensor NLRs form well-supported phylogenetic clusters (Wu *et al*. 2017), we aimed to identify NLRs belonging to either group using a comparative phylogenetic clustering analysis with functionally characterized helper or sensor NLRs. We observed the formation of two phylogenetically distinct sweetpotato NRC-H subclades from the NRC0/1 references. Consistent with our observations, Adachi *et al*. (2023) identified the formation of two small subclades apart from the NCR1/2/3, NRC4, and NRCX clades in a phylogenetic analysis that included CNLs from *Arabidopsis*, sugar beet, tomato and *N. benthamiana*. Helper NLRs are critical hubs in the NLR network (Białas *et al*. 2018). We documented the expansion of *Ipomoea* NRC sensors and provided an example for an NRC superclade experiencing diversification in the NRC-S subclades. Our clustering analysis aids in identification of candidate NRC proteins that can be assessed for importance within the sweetpotato NLR network via functional studies (Kourelis *et al*. 2022).

The domesticated sweetpotato NLRome contained more than 1,000 NLRs for some genotypes while its wild ancestor harbored roughly half in our study. Notably, other clonally propagated polyploid crops harbor unusually large NLRomes with sizeable expansions (Jia *et al*. 2015; Tang *et al*. 2022). Such expansion of NLRs in apples is hypothesized to be a result of domestication (Jia *et al*. 2015). Tang *et al*. (2022) postulated that the potato NLRome expansion may have co-evolved with the emergence of clonal propagation approximately 7.3 million years ago. Our observations in cultivated sweetpotato and wild relative genotypes support this hypothesis. While domestication secured an adaptable and staple sweetpotato crop, the functional significance of harboring a large NLRome remains unknown but worthy of exploration. By conducting a presence/absence analysis of orthogroups across genotypes, we confirmed that a third of orthogroups belonged to 10 or more genotypes, however, they accounted for two thirds of the shared NLR counts. Only 5.9% of NLRs were uniquely observed within a single genotype with a high proportion of NLRs shared across two or more genotypes reflecting true conservation. This pattern of conservation in sweetpotato contrasts with both *A. thaliana* and Solanaceae pan-NLRome analyses that reported a limited core of NLRs across genotypes accounting for only a small proportion of all NLR genes (Van de Weyer *et al*. 2019; Seong *et al*. 2020; Barragan and Weigel, 2021).

Detection of invading pathogens and activation of immune response represents a major role of NLRs in plants (Jones *et al*. 2016). Largely influenced by pathogen evolution, NLRs exhibit patterns of rapid and dynamic evolution at the intraspecific level (Lee and Chae, 2020). However, we lack knowledge on the level of divergence occurring among sweetpotato genotypes and their wild relatives. Time calibrated phylogenies indicate that the hexaploid sweetpotato likely diverged from its closest wild relative, *I. trifida*, over 1 million years ago with a significant portion of sweetpotato diversity largely predating the onset of agriculture (Muñoz-Rodríguez *et al*. 2018, 2019). Equipped with our curated CNL annotations, we examined the phylogenetic distance among CNLs of sweetpotato genotypes and its ancestor, *I. trifida*. Our analysis revealed that CNLs remain largely conserved within *I. batatas* genotypes and *I. trifida*, with only a few CNLs diverging. This supports our OG analysis, which suggested a largely common NLRome among *I. batatas* genotypes. We postulate that an ancient wild relative likely provided the NLRs that remain common across the majority of *I. batatas* genotypes indicating conservation of the sweetpotato gene pool over domestication, clonal propagation, and breeding. In a meta-analysis that included barley, cucurbits, and sunflower NLRs, Baggs *et al*. (2017) postulate that NLR count variation between wild relatives and cultivated genotypes is a consequence of domestication bottlenecks. In our study, we observed the opposite and hypothesize that polyploidization and outcrossing breeding enriched NLRs in sweetpotato. Our gene models were predicted using long read cDNA evidence suggesting transcriptionally active NLR genes. However, some NLR genes may be epigenetically regulated (Lai and Eulgem, 2018) and/or constitute the large functional redundancy required by the helper/sensor model proposed by Wu *et al*. (2018). In addition, we discovered a small set of sweetpotato CNLs with notable patristic distance from the 3 diploid wild relatives. These CNLs represent excellent candidates to measure directional selection as they appear to be an important group of NLRs undergoing divergence among some sweetpotato genotypes. Elucidating if the divergence observed among some genotypes arose as modulation of inappropriate activation of NLR defense signaling by surrounding microbes/pathogens in the environment could help identify NLRs acting as susceptibility genes (Warmerdam *et al*. 2018). Perhaps, these CNLs activation becomes detrimental for some genotypes if over-triggered, equating to energy loss or pathogen recognition fatigue (Karasov *et al*. 2017).

Sweetpotato exhibit self-incompatibility, which results in high levels of heterozygosity (Wu *et al*. 2018b). A recent study by Seong *et al*. (2022) suggested that for highly heterozygous plant species, RenSeq fails to resolve the complexity of NLR diversity. Judging by our number of gene models that correspond to truncated/partial NLRs (i.e. NLs, CNs, TNs), we agree with their assertion. However, we hypothesize that for highly heterozygous hosts a more stringent cutoff for complete NLR loci results in a highly accurate set of NLRs. We conclude this based on the proportion of NLR counts in the diploid wild relative species included in our study. Focusing on a set of complete NLR domains allowed us to obtain relatively high mapping rates (84%) of our RenSeq data on the reference genome assemblies. Thus, we postulate that a higher level of filtering can be applied to the gene models that contain canonical sets. We expected that some models would be miss-annotated, but by focusing on models that are likely accurate based on domain architecture and consistency across genotypes, we were able to improve our knowledge of NLR diversity in heterozygous sweetpotato. Partial NLR models excluded from this study may represent misannotated NLRs that could be prioritized for manual curation in future studies in the same manner as Seong et al. (2022), Lin *et al*. (2022), and Van de Weyer *et al*. (2021) implemented for *Solanum* wild relatives and *Arabidopsis* ecotypes.

Capturing NLRs using RenSeq facilitates anchoring NLR contigs to reference genomes (Jupe *et al*. 2013; Arora *et al*. 2019). Here, we used RenSeq derived contigs to identify regions in the chromosome scale assembly harboring NLR loci, focusing only on NLR loci predicted to contain full-length NLRs. We observed heavy NLR clustering and separation by canonical domain types with some CNL and TNL loci restricted to specific chromosomes. Anchoring NLR contigs to chromosomes remains an intricate task that relies on the quality of sequences flanking the NLR loci which largely depends on the length of the original molecules captured during the enrichment step (Barragan and Weigel, 2021). A limitation of our approach is that there may be other NLR loci with duplication events or lower mapping identity that we excluded but may represent viable breeding targets. However, our NLR loci anchoring analysis allowed us to map the vast majority of the full-length NLR loci representing a nearly complete map of NLRs in the hexaploid sweetpotato. The NLR coordinates and clustering that we documented among chromosomes could facilitate breeding efforts by improving resolution for QTLs of interest. Future work leveraging synteny analyses and phylogenetic reconstruction could explore the evolutionary dynamics of NLR loci across the different sub-genomes, particularly in relation to wild relatives such as *I. trifida*. Our RenSeq panel included parents of mapping populations segregating for resistance to different diseases. Subsequent RenSeq studies in combination with bulked segregant analysis (BSA) may help identify NLRs conferring resistance to particular pathogens (Lin *et al*. 2022). As a first step towards that goal, we conducted RenSeq in a polyploid highly heterozygous staple crop, potentially contributing to NLR cataloging efforts in orphan crops (Ye and Fan, 2021).

In conclusion, we described the NLRome of hexaploid sweetpotato and its wild relatives, identifying a highly conserved NLR catalog among sweetpotato genotypes. Our annotation and phylogenetic analysis reveal an expanding CNL clade with potential sensor and helper NLRs to functionally characterize. We recorded low divergence between *I. batatas* and *I. trifida* CNLs also suggesting a conserved NLRome. This study provided the nearly complete NLR loci coordinates within the sweetpotato chromosome level assembly. Our RenSeq study provides a catalog of NLR genes that will accelerate breeding for disease resistance and improve our understanding of the evolutionary dynamics of NLRs in sweetpotato.

## ACKNOWLEDGMENTS

We thank Dr. Zhangjun Fei for pre-publication access to the hexaploid genome assembly that we used for NLR anchoring and clustering analysis. We want to thank Chris Heim for assistance on maintenance of sweetpotato genotypes in the greenhouse. We specially thank Kamil Witek and Brian Brunelle for helpful discussions on the implementation of RenSeq in our study. We also thank Lisa Lowe for her assistance with parallel optimization of NLR assembly and phylogeny software. We acknowledge the computing resources provided by NC State University High Performance Computing Services Core Facility (RRID:SCR_022168). We also thank Lindsey Becker for commenting on an earlier draft of the manuscript. The Vegetable Pathology Lab at NC State University for their kind support, especially Katie Rose Ketzes and Hunter Collins. This work was supported by the United States Department of Agriculture (USDA), National Institute of Food and Agriculture (NIFA) Award 2168-207-2023550, the Bill and Melinda Gates Foundation (INV-002971), the Foundation for Food and Agriculture Research (FFAR) Fellowship Program, the NC Sweetpotato Commission, and the NC State Hatch Project NC02890.

## AUTHOR CONTRIBUTIONS

CHPR and LMQO designed the project. CHPR designed the bait library. CA, GCY, and KP provided plant material. CHPR performed DNA extractions. CHPR, AS, DB, and MFS performed RenSeq library construction. CHPR and KLC conducted assembly and structural annotation. CRB and MK provided long read transcript evidence for gene annotation. CHPR, KC, GCC, analyzed the data and results. CHPR wrote the manuscript.

---

**Figure S2 Sweetpotato and wild relative genomes harbor a diverse catalog of NLRs.** NLRs grouped in the canonical domains TNL, CNL, and RNL domain architecture and non canonical BNLs and NLR-IDs. Counts for each genotype are plotted as black circles and densities shown as half violin plots. Domain architecture and abbreviations are as shown in Figure S1.

