## Supplementary material for "A reference-quality NLRome for the hexaploid sweetpotato and diploid wild relatives": Methods S1

**Plant material, growth conditions and DNA extractions.** To capture the global and local diversity of *Ipomoea batatas,* we included 32 hexaploid *I. batatas* genotypes and three diploid wild *Ipomoea* sp. genotypes. We selected a set of 32 representative *I. batatas* genotypes based on their importance and potential as parents of mapping populations. This *I. batatas* panel included land races, cultivated, and advanced breeding lines. We also included three wild *Ipomoea* species including *I. littoralis* (PI 573335), *I. triloba* (NCNSP0323), and *I. trifida* (NCNSP0306); the former is considered the progenitor of cultivated sweetpotato (Table S5). We included 32 hexaploid *I. batatas* genotypes and three diploid wild *Ipomoea* sp. genotypes. Our sweetpotato hexaploid panel included a set of 10 resistant and 10 susceptible genotypes for both *Meloidogyne enterolobii* and *Fusarium solani* (Yencho *unpublished*) (Table S5). Mother plants were maintained in a NC State University greenhouse under a 16 h-light/8 h-dark photoperiod and grown in 6 in pots filled with peat moss-vermiculite potting media (Conrad Fafard Inc. FL, USA). Plants were watered twice daily and fertilized with 20N–10P–20K (J.R. Peters Inc. PA, USA) every week. Certified pathogen-free slips (cuttings) from contemporary and commercial genotypes were obtained from the Sweetpotato Micropropagation Unit at NC State University (NC State) and the NC State sweetpotato breeding program (https://potatoes.cals.ncsu.edu/). A historical *I. batatas* genotype and three wild relatives (*I. triloba, I. trifida* and *I. littoralis*) were obtained from the USDA Germplasm Resources Information Network (GRIN). Genomic DNA of young leaf tissues obtained from one to two plants per genotype was extracted using NucleoBond HMW DNA Kit (MACHEREY - NAGEL Inc., PA, USA). We extracted approximately 10 µg of genomic DNA per genotype to allow downstream library preparation. To confirm high molecular weight and DNA quality for RenSeq library preparation and long read sequencing, we performed size/integrity analysis and quantification of nucleic acids in an Agilent 2200 TapeStation (Agilent Technologies, Inc., CA, USA).

**NLR gene enrichment sequencing.** To design our target NLR bait library, we used NLR-parser, a benchmarked NLR annotation tool, that provides sequence coordinates of complete and partial NLRs in a set of query sequences (Steuernagel *et al*., 2015). We scanned the available genomic resources for sweetpotato, including two diploid wild relatives (*I. trifida* and *I. triloba*) high-confidence coding DNA sequences (cDNA), a transcriptome assembly from the hexaploid genotype Beauregard, and cDNA from *I. nil* (Steuernagel *et al.* 2015). The cDNA sequences represent spliced transcript models, including untranslated regions, and were chosen to ensure comprehensive representation of gene models. Only complete putative NLR sequences were used to create a bait library as described by Jupe et al. (2013). In brief, the library was composed of 120-mer biotinylated RNA baits starting from the first nucleotide following the predicted coding region. Sequences were tiled at least 2 times with a 60 nucleotide overlap and bait sequences containing N residues were subsequently removed. Bait candidates were filtered against plastid sequences. The resulting 38,694 baits were synthesized by Arbor Biosciences (Arbor Biosciences, Ann Arbor, MI, USA). A total of 10 µg of high molecular weight (HMW) genomic DNA (gDNA) from each genotype were fragmented with a sonicator (Covaris Inc., Woburn, MA, USA). Sheared gDNA was size-selected to obtain 3-5 Kb fragments with blue pippin BLF7510 cassettes (Sage Science, Beverly, MA, USA). NLR capture followed MYbaits v4.0 protocol with the following modifications. A total of 1.5 µg of size selected DNA was hybridized with the baits using the total reaction volume suggested. For each reaction, the hybridization master mix was prepared to include: 9.25 µl of Hyb N, 3.5 µl of Hyb D, 0.5 µl of Hyb S, 1.25 µl of Hyb R, 5 µl of SeqCAP (Roche, Indianapolis, IN), 0.5 µl of Block A, and 5.5 µl of bait library. For the capture reaction, the final reaction volume was 30 µl which included 23 µl of hybridization mix and 7 µl of size-selected gDNA. The cycling conditions for hybridization included a 10 min denaturation step at 95°C followed by a touchdown phase that reduces the temperature 0.1°C per second until reaching 65°C, and maintained at 65°C for 16 to 24 hours. The enriched libraries were recovered by adding 50 µl of washed Dynabeads MyOne Streptavidin C1 beads and warmed up to 65°C as suggested by the manufacturer. Binding and elution of captured gDNA fragments were accomplished as suggested in MYbaits v4.0 protocol. To test the enrichment efficiency, we performed qPCRs on four NLR targets included in the bait library and Actin gene (3 technical replicates for each capture library). Primers used for qPCR are listed in Table S6. Quantitative PCRs (qPCRs) were performed for a total 20 µl volume using 10 µl of 2X SYBR Green Mix (Thermo Fisher Scientific, Rochester, NY),4 µl of diluted RenSeq library (0.25ng/ µl), and 0.5 µl of each primer (10 µM). qPCR was run on the CFX96 Real-Time System C1000 thermal cycler (BioRad, Hercules, CA) using the following program profile: (1) 50°C, 2 min, (2) 95°C, 2 min; (3) [95°C, 15 sec, then 60°C, 1 min] x 40, 65°C, 0.05 min followed by a temperature gradient from 65°C to 95°C. The Ct value difference between before and after-capture libraries were quantified. Captured libraries that passed the enrichment efficiency check (8 -10 cycle difference) were amplified using high fidelity KAPA enzyme 1 U/µL (Roche, Indianapolis, IN) and subsequently prepared for PacBio SMRT sequencing following the NC State-GSL (Genomic Sciences Laboratory) standard recommendations for 4-10 kb library preparation. The libraries were sequenced using the Sequel PacBio platform at the NC State GSL.

**Assembly, structural and functional NLR annotation.** Circular Consensus Sequencing (CCS) reads were generated from raw subreads using *ccs* with three full passes and 90% accuracy (version 6.4.0; -j 16 --min-rq 0.9 --min-passes 3 --max-length 50000) (PacBio, 2022). Each set of CCS reads from each genotype was processed to remove the adapters and barcodes using Cutadapt (version 1.16; -u 65 -u -65 -e 0.05 -m 150) (Martin, 2011). Trimmed reads were searched for chimeric reads using BLASR and removed manually (version 5.3.5; -m 1 --bestn 10) (Chaisson & Tesler, 2012). To assess the number of CCS reads containing at least one NLR bait sequence, we conducted a BLAST searched of the entire NLR bait library on each genotype CCS read library using BLAST+ (v.2.9.0; -num_threads 8 -max_target_seqs 1 -max_hsps 1 -outfmt 6). The resulting hit table was parsed to determine the number of reads with > 80% identity over a stretch of > 96 bp as recommended by Giolai et al. (2016). We also calculated the number of reads containing NLR motifs using NLR-parser that (unlike NLR-tracker) can scan DNA sequences. We reported the number of reads containing NLRs as defined by NLRparser (Steuernagel *et al.*, 2015). Only non-chimeric reads were assembled using Canu (version 1.6; genomeSize = 12.5m correctedErrorRate = 0.010 -minOverlapLength = 350 -trimReadsCoverage = 1 -minReadLength = 1000) (Koren *et al.*, 2017). Contig assemblies were assessed for performance by searching the contig sequences with NLR-annotator, a benchmarked tool to detect NLR motifs in contig sequences (Zhang, 2020). We reported the number of NLRs containing complete NLR motifs as defined by NLR-annotator and the number of contigs containing at least one NLR locus per read. A custom MAKER2 annotation pipeline was designed to predict NLR gene models (version 2.31.9; pred_flank = 200, keep_preds = 1, split_hit = 10000, tries = 2, AED_threshold=1) (Holt & Yandell, 2011). Protein evidence from *I. trifida, I. triloba* and *I. nil* was externally aligned using Exonerate (version 2.2.0; --model protein2genome --bestn 5 --minintron 10 --maxintron 3000). Transcript evidence consisted of Beauregard ONT (Oxford Nanopore Technology) full-length cDNAreads generated from combined leaf, fibrous, and storage root tissues cDNA (Buell *unpublished*). Trimmed, oriented, and filtered Beauregard cDNA reads were mapped to the RenSeq contigs of each genotype using Minimap2 (version 2.17; -N 1 -t 20 -ax splice -g2000 -G5k) (Li, 2018). The resulting BAM format files were assembled into transcripts using StringTie2 (version 2.0; -m 300 -t -c 2.5 -f 0.05 -g 50 -p 30) (Kovaka *et al.*, 2019). The resulting GTF format files from both protein and transcript evidence were sorted and converted to GFF3 format to input into MAKER2*.* A first round of MAKER2 was performed to infer gene predictions directly using the est2genome parameter while providing both protein and transcript evidence. Gene predictions from representative hexaploid (Beauregard) and diploid (*I. trifida)* were used to train AUGUSTUS (Korf, 2004) and SNAP HMMs (Hidden Markov Models) (Stanke *et al.*, 2008). Beauregard and *I. trifida* hmms were then used for final gene prediction in *I. batatas* genotypes and *Ipomoea* wild relatives respectively. The second round of MAKER2 used both protein and transcript evidence and the respective trained HMMs to generate the final gene predictions. For each genotype, a MAKER2 standard gene set was produced that consisted of gene model predictions that had evidence support or that contained a Pfam domain. Our MAKER2 pipeline omitted the customary repeat masking step as it tends to shorten the gene model and misannotate the LRR domain in NLRs for repeat sequences (Bayer *et al.*, 2018). Evidence and gene model inspection was carried out in the Integrative Genomics Viewer software (IGV)(version IGV 2.14.x) (Robinson *et al.*, 2011). NLR protein models were classified based on their multi-domain architecture using the benchmarked NLR-tracker tool (Kourelis *et al.*, 2021). We followed the NLR-tracker definition of NLRs, which defines NLRs as those protein models that contain an NB-ARC domain and at least one additional domain. However, this definition allows for many truncated or partial NLRs to be included and therefore we arbitrarily set a threshold to select complete NLR protein models that consisted of: models carrying the canonical NB-ARC, LRR, and one the N or C-terminal domains (CC, TIR, RPW8, B, CID-J, and IDs) (Figure S1). The current notion of integrated domains suggest that NLRs carrying IDs represent key pathogen effector targets, therefore we looked at their distribution in our dataset by grouping them into 6 distinct subclasses: ONL, OCNL, CNLO, OTNL, TNLO, and RNLO (Figure S1) (Cesari *et al.*, 2014). To allow future reference of the annotated NLR models, we named the FASTA and GFF files as follows: iba_apa00003g02.1; in which “iba” stands for *Ipomoea batatas*; “apa” corresponds to the genotype Apache; the following 5 digits identifies the RenSeq contig number where the gene model was predicted; “g0#” specifies the gene number in the respective contig i.e. g02 means this is the second gene in the contig; finally, “.1” indicates that this gene has a corresponding transcript evidence associated in the transcript FASTA file. Table S4 lists the general abbreviation/naming convention for each of the 35 genotypes included in this study. Each FASTA header and the GFF3 file includes the simplified domain architecture (ea. “CNLO”) as a note.

**Comparison of NLR content in standard genome annotation.** To highlight the differences in NLR annotation outcomes between standard genome annotation projects and RenSeq/NLR tailored annotation pipelines, the proteomes from 8 plant species were downloaded from individual genome project repositories. This panel included the proteomes of recently available *Ipomoea batatas* (cv. Beauregard, representative high confidence gene model set version 1) (*pre-publication version*), the proteomes of two diploid sweetpotato wild relatives *I. trifida* (NSP306, representative high confidence gene model set version 3) and *I. triloba* (NSP323, representative high confidence gene model set version 3) (Wu *et al.*, 2018b), the proteomes of 4 Solanaceae species including *Solanum tuberosum* (Doubled monoploid potato DM 1-3 516 R44, High Confidence Gene Model Set - v6.1) (Pham *et al.*, 2020), *Solanum lycopersicum* (cv. Heinz 1706; annotation release ITAG4.1) (Hosmani *et al.*, 2019), *Capsicum annuum* (cv. CM334 (Criollo de Morelos 334); annotation version 1.55) (Hulse-Kemp *et al.*, 2018), and *Nicotiana benthamiana* (Accession Nb-1, annotation version 2.6.1) (Bombarely *et al.*, 2012). In addition, we included as an outgroup the proteome of the Brassica *Arabidopsis thaliana* (Ecotype Col-0; Araport11 annotation) (TAIR, 2022). All protein sequences were scanned through NLR-tracker and categorized as NLRs if the sequence contained an NB-ARC domain and at least 1 additional domain. We compared each genome annotation NLR count with that reported for the corresponding RenSeq projects for the same plant species. The NLR counts from RenSeq projects were collected as reported in each RenSeq study.

**NLR phylogenetic diversity in 6X and 2X *Ipomoea* spp.** To explore the diversity of NLRs in the 32 *I. batatas* hexaploid and 3 diploid wild relatives, we constructed NLR phylogenies for complete NLRs only. We employed deduplicated amino acid sequences of the NB-ARC domains extracted by NLR-tracker to produce a multiple sequence alignment (MSA). We included a diverse panel of 35 refPlantNLRs that encompasses major functionally annotated and phylogenetically diverse NLRs (Kourelis et al., 2021). The MSA was generated using the globalpair alignment in MAFFT (version 7.490; --large --globalpair) (Rozewicki *et al.*, 2019). After visual inspection and gap threshold testing, the MSA was trimmed to remove any columns containing gaps in 80% or more of the sequence alignment using Trimal (version 1.4.1; -gt 0.2), which ensured the inclusion of shorter NB-ARC reference sequences (Capella-Gutiérrez *et al.*, 2009). A maximum-likelihood tree was inferred from the resulting MSA of 379 columns and 29,553 sequences using ExaML under the per-site rate category model (version 3.0.17; -m PSR) (Kozlov *et al.*, 2015). We inferred 6 randomized stepwise addition order parsimony-based starting trees required for ExaML using RAxML (version 8.1.20; -y -N 6 -m PROTCATJTT -p 12345) (Stamatakis, 2014). The final tree was visualized in the Interactive Tree Of Life (iTOL) software (version 6.6) and arbitrarily rooted on the branch connecting TNLs and non-TNL clades (Letunic & Bork, 2007). To provide additional context on potential candidate NLRs for functional studies in sweetpotato, we pinpointed the phylogenetic position of the NRC superclade in the large sweetpotato NLR phylogeny. We pruned and extracted NLRs that clustered with functionally characterized NRC0, and NRC1 and visualized the phylogeny in iTOL.

**Phylogenetic distance analysis.** To evaluate NLR conservation between diploid wild relatives (N = 3) and *I. batatas* genotypes (N= 32), we calculated phylogenetic distance among complete CNLs as they represent the largest expanding NLR clade in our study. First, we extracted NB-ARC deduplicated domains corresponding to complete CNLs in all 35 genotypes. A total of 6 NB-ARC datasets were generated; three datasets represented the 3 wild relatives individually (*I. trifida*, *I. triloba*, and *I. littoralis*) and the other 3 datasets included all 32 hexaploid *I. batatas* genotypes in combination with a single wild relative (32 + Itf; 32 + Itb; 32 + lito). We aligned the NB-ARC amino acid sequences using MAFFT (version 7.505; --large --globalpair) and removed columns containing gaps in 80% or more of the sequence alignment using Trimal (version 1.4.1; -gt 0.2). We generated 3 NLR phylogenies with the combined 32 *I. batatas* and each wild relative, *I. trifida*, *I. triloba*, and *I. littoralis*, respectively. Three individual diploid wild relative phylogenies were generated for *I. trifida*, *I. triloba*, and *I. littoralis.* We calculated the phylogenetic (patristic) distance between each pair of CNLs in the diploid wild relatives and their corresponding closest CNLs in the 32 hexaploid sweetpotato genotypes. We used the publicly available python script *Phylogenetic distance analysis* (https://github.com/slt666666/Phylogenetic_distance_plot2) to calculate the distance table and visualized distance against each diploid wild relative phylogeny in R using ggplot2.

**Orthology inference, refinement, and classification.** We inferred orthologous groups using a pairwise global amino acid similarity approach over the length of the NB-ARC domains extracted by NLR-tracker from each of the 35 genotypes. First, we produced an all-against-all global local alignment in BLAST+ (v.2.9.0; -evalue 1e-20). Second, using the results of the BLAST search, we initialized a distance matrix for all 97,505 NB-ARC amino acid sequences from all 35 genotypes. For any pair of such genes with a BLAST Evalue > 1e−20, we used the global alignment algorithm of Needleman and Wunsch (1970) to align the two sequences: we then computed the pairwise amino acid percent difference between the two sequences and stored this value in the matrix. We created a graph where edges connected nodes (sequences) with percent identity >98.5%. We then inferred orthogroups to be connected components within this graph. We explored a range of cutoffs values between 95% and 99.25%. A cutoff of 98.5% conferred the largest number of orthogroups with between 15 and 55 members. We refined the orthogroups by assigning the predicted NLR domain architecture to each sequence belonging to each orthogroup. We kept orthogroups that contained sequences assigned to complete NLR domains (RNLO, TNLO, CNLO, OCNL, OTNL, ONL, BNL, BCNL, CNL, TNL, RNL, CCNL, and TNLJ) (Figure S1). The final set of refined orthogroups was classified into each of the complete NLR domain architectures. Manual annotation was required for 320 orthogroups, who exhibited more than one NLR domain architecture. In these cases, we assigned the orthogroup to the longest domain architecture (i.e. CNL vs CNLO, we kept CNLO). The orthogroup counts per genotype were converted into a presence/absence matrix to examine orthogroup distribution among 32 sweetpotato genotypes and assign cloud, shell, and core categories. We classified orthogroups into the cloud category if the orthogroups were shared by < 10 genotypes, the shell category included orthogroups shared between 11 and 20 genotypes, finally the core orthogroup category included orthogroups shared by > 21 genotypes. The co-occurrence of orthogroups and their corresponding category across genotypes was visualized in R using ggplot2.

**Genomic anchoring of complete NLRs in hexaploid Beauregard.** To examine the chromosome level clustering and location of NLRs in hexaploid sweetpotato, we anchored the hexaploid Beauregard RenSeq contigs into the recently haplotype-resolved chromosome-scale Beauregard genome assembly (*pre-publication version*). Raw RenSeq contigs were filtered using the NLR-tracker complete NLR domain architecture predictions that match the contig and gene header in the FASTA file. We mapped the resulting contigs to the hexaploid chromosomes using Minimap2 (version 2.17; -N 1 -t 15 -ax asm5 -g2000 -G5k). The BAM format alignment file was sorted, indexed and converted into a SAM format using SAMTools (version 1.9). We inspected the contig alignment in Geneious Prime (version 2022.0.2), removed secondary and supplementary alignments in both strands, and kept alignments with quality tag = 60. We filtered further the alignment and removed any contigs mapping with an identity threshold below 99%. This filtering approach allowed us to remove contigs that mapped more than once in any position in the genome. We visualized NLR contigs along chromosomes using the R package RIdeogram (version 0.2.2) (Hao *et al.*, 2020). We repeated this process for the *I. trifida* RenSeq contigs and its corresponding genome assembly (Wu *et al.*, 2018b).
